# Boreal river impoundments caused little change in fish diversity but clear community assemblage shifts: A multi-scale analysis

**DOI:** 10.1101/129403

**Authors:** Katrine Turgeon, Christian Turpin, Irene Gregory-Eaves

## Abstract

1. Hydroelectricity is often presented as a clean and renewable energy source, but river flow regulation and fragmentation caused by dams are recognized to impact aquatic biodiversity in temperate and tropical ecosystems. However, the effects of boreal river impoundment are not clear as the few studies that exist have not been able to separate the hydrological changes brought about by dams from other factors (*e.g.* fish stocking, and species introduction).
2. We adopted a multi-scale analysis to examine changes in nearshore fish communities over 20 years (spanning before and after impoundment) using a network of 24 sampling stations spread across from four reservoirs and two hydroelectricity complexes located in the boreal region (Northern Québec, Canada). Given the remote location, confounding factors were minimal.
3. We found no strong temporal trends in alpha- and gamma-diversity in impacted stations (upstream and downstream of the dam) relative to reference sites across the three spatial scales. Using beta-diversity analyses, we also detected a high stability in fish composition over time and space at the complex and reservoir scales.
4. At the scale of the sampling stations, we observed higher rates of species turnover (beta-diversity) coincident with the time of reservoir filling and shortly after. Likewise, we detected species assemblage shifts that correlated with time since impoundment only at the sampling station scale. This pattern was masked at the complex and reservoir scales.
5. *Synthesis and applications*. Overall, the isolated effect of impoundment in these remote boreal ecosystems caused no loss of species and little change in fish diversity over 20 years, but resulted in substantial species assemblage shifts. Our work shows that examining community data at different scales is key to understand the anthropogenic impacts on fish biodiversity.

## 1. Introduction

In response to increased demand for energy, many large dams are currently in operation, or are being constructed to provide hydroelectricity (Grill *et al.* 2015; Winemiller *et al.* 2016). Dams transform large rivers into large reservoirs, affecting numerous important physical, chemical and biological processes (Ward & Stanford 1995; Friedl & Wüest 2002). Dams also fragment rivers by creating barriers to movement (Nilsson *et al.* 2005; Pelicice, Pompeu & Agostinho 2015), and alter the natural hydrological regime of the ecosystem (*i.e.*, discharge and water levels) upstream and downstream of the dam (Kroger 1973; Poff *et al.* 2007). These modifications are susceptible to affect the overall biodiversity and ecosystem functions (Rosenberg, McCully & Pringle 2000; Vörösmarty *et al.* 2010; Liermann *et al.* 2012).

The effects of impoundment on fish communities have been extensively studied in temperate (Martinez *et al.* 1994; Bonner & Wilde 2000; Gido, Matthews & Wolfinbarger 2000; Taylor, Knouft & Hiland 2001; Gehrke, Gilligan & Barwick 2002; Quinn & Kwak 2003), and more recently in tropical ecosystems where many new dams have been constructed (de Mérona, Vigouroux & Tejerina-Garro 2005; Agostinho, Pelicice & Gomes 2008; Li, Madden & Xu 2012; Lima *et al.* 2016). However, very little is known about the effects of impoundment in boreal ecosystems (but see Tereshchenko & Strel’nikov 1997; Sutela & Vehanen 2008). This deficiency is surprising considering that hydroelectricity is a major source of energy in some Nordic countries (*e.g.*, Norway: 96% of domestic electricity generation, Iceland: 70%, Canada: 58% and province of Québec in Canada: 95%; IEA 2016).

Long-term monitoring of fish assemblages in reservoirs is critical (Elliott 1990; Gido, Matthews & Wolfinbarger 2000) because reservoirs are young (average of < 60 years), novel ecosystems, and they are highly dynamic in the first decades following impoundment (*i.e.*, non-trophic equilibrium phase, Grimard & Jones 1982; Turgeon *et al.* 2016). Moreover, the time needed for the fish community to adapt (or not) to the new reservoir conditions, will depend on several factors such as geographic location, reservoirs characteristics (*i.e.*, reservoir area, water quality), dam operation and management (*i.e.*, drawdown), complexity of the food web, and species life history traits. These potential sources of variability stress for the importance of replication. To extract generalities about the effects of impoundment on fish community across all latitudes, we need to improve our approach by having an exhaustive examination of the following elements: 1) observations on fish communities spanning before to after impoundment, collected routinely for many years, 2) data collected from multiple sampling stations downstream and upstream of the dam and 3) parallel measurements made in reference sites to identify climatic or other regional environmental drivers of change.

In this study, we took a multi-scale approach to examine how the impoundment of rivers affects fish communities in four boreal reservoirs using a long-term dataset collected by Hydro-Québec (Fig. 1). This dataset consists of a large network of 24 sampling stations (including upstream and downstream stations as well as reference sites) and spans from before the construction to 10 or 20 years after the start of its operation, allowing for one of the most thorough and robust evaluations of how impoundment affect fish communities. Moreover, because of its remoteness, this dataset provides a rare opportunity to isolate the effect of impoundment on fish communities from other anthropogenic confounding factors.

**Figure 1.**
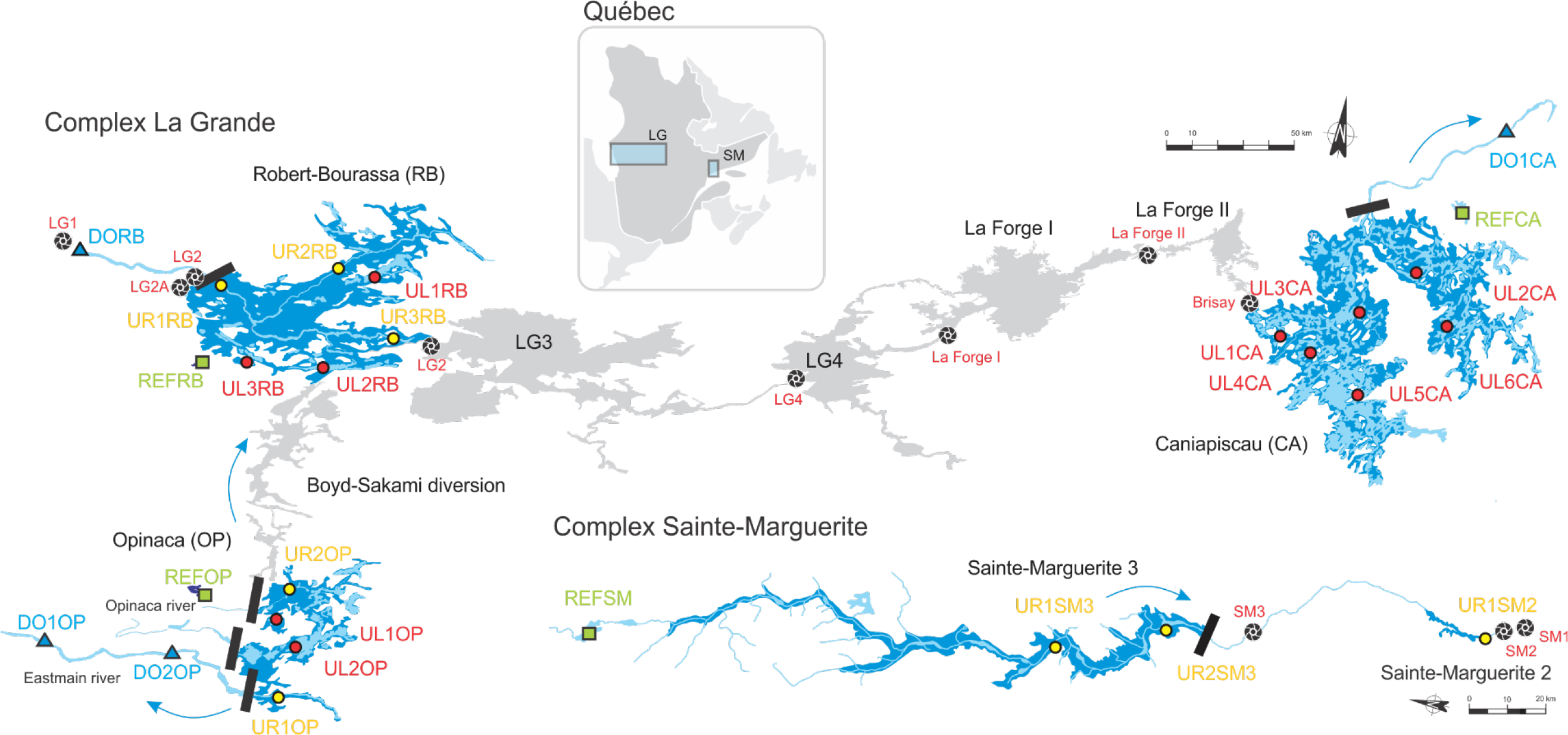
Map representing the before (light blue) and after (dark blue) impoundment hydrological conditions, and the location of the sampling stations in the La Grande hydroelectricity complex from three reservoirs; RB, OP and CA and 4 sampling stations in Sainte-Marguerite complex, Northern Québec. Stations located upstream of the dams that were in a river before impoundment are represented by yellow circles, and the ones that were in lakes before impoundment are represented by red circles. Sampling stations that were located downstream of the dams are represented by a blue triangle, and reference sites paired with each reservoir are represented by green squares. Dams are represented by a black line and power station by a turbine symbol. Reservoirs that are not the focus of our study but in the region, (LG3, LG4, LaForge I & II) are presented and were impounded at the following dates: 1981, 1983, 1993 and 1983 respectively.

## 2. Materials and Methods

### 2.1 Study sites

#### LG complex

The La Grande Rivière hydroelectric complex (hereafter called LG complex) is located on the eastern side of James Bay (Québec, Canada), on the Canadian Shield. The LG complex resulted in the creation of seven large reservoirs (Table 1, Fig. 1), and in the diversion of three large rivers, the Caniapiscau (water flow at its mouth reduced by 43%), the Eastmain (reduction of 86%) and the Opinaca (reduction of 86%; Roy & Messier 1989). Through these hydrological changes, the average annual discharge in La Grande Rivière has increased from 1700 m^3^•s^-1^ to 3400 m^3^•s^-1^ (Roy & Messier 1989). Data have been routinely collected from three reservoirs in the LG complex: Robert-Bourassa (RB; impounded in 1979), Opinaca (OP; impounded in 1980) and Caniapiscau (CA; impounded in 1982; Table 1, Fig. 1). Reservoirs LG3, LG4 and LaForge II have been impounded in 1981, 1983 and 1983 respectively. Laforge I have been impounded later in 1993 (Fig. 1). The territory is free of other industrial activities and sparsely occupied by the Indigenous Cree peoples. Some mitigation measures over the years included fish habitat improvement (new spawning area, vegetation control, creation of shelters, containment) and the maintenance of fish movement (*i.e.*, migratory pass in Robert-Bourassa was created in 1980). Each of the three reservoirs was paired with a natural lake in proximity to the reservoir (“REF” stations; Fig. 1). Fish community data were collected in stations downstream of the dam (“DO” stations; Fig. 1) and upstream of the dam (“UR” if the station was a river or stream before and “UL” if the station was a lake before impoundment; Fig. 1). We expected that UR stations would demonstrate a more pronounced change in diversity and fish assemblages than UL stations because of the drastic change in habitat from lotic to lentic conditions. One downstream station (DORB) had an increased flow after impoundment, and the three others (DO1OP, DO2OP and DO1CA; Fig. 1) had decreased flow because sampling stations were in rivers that were diverted to create the reservoirs.

**Tabel 1.**
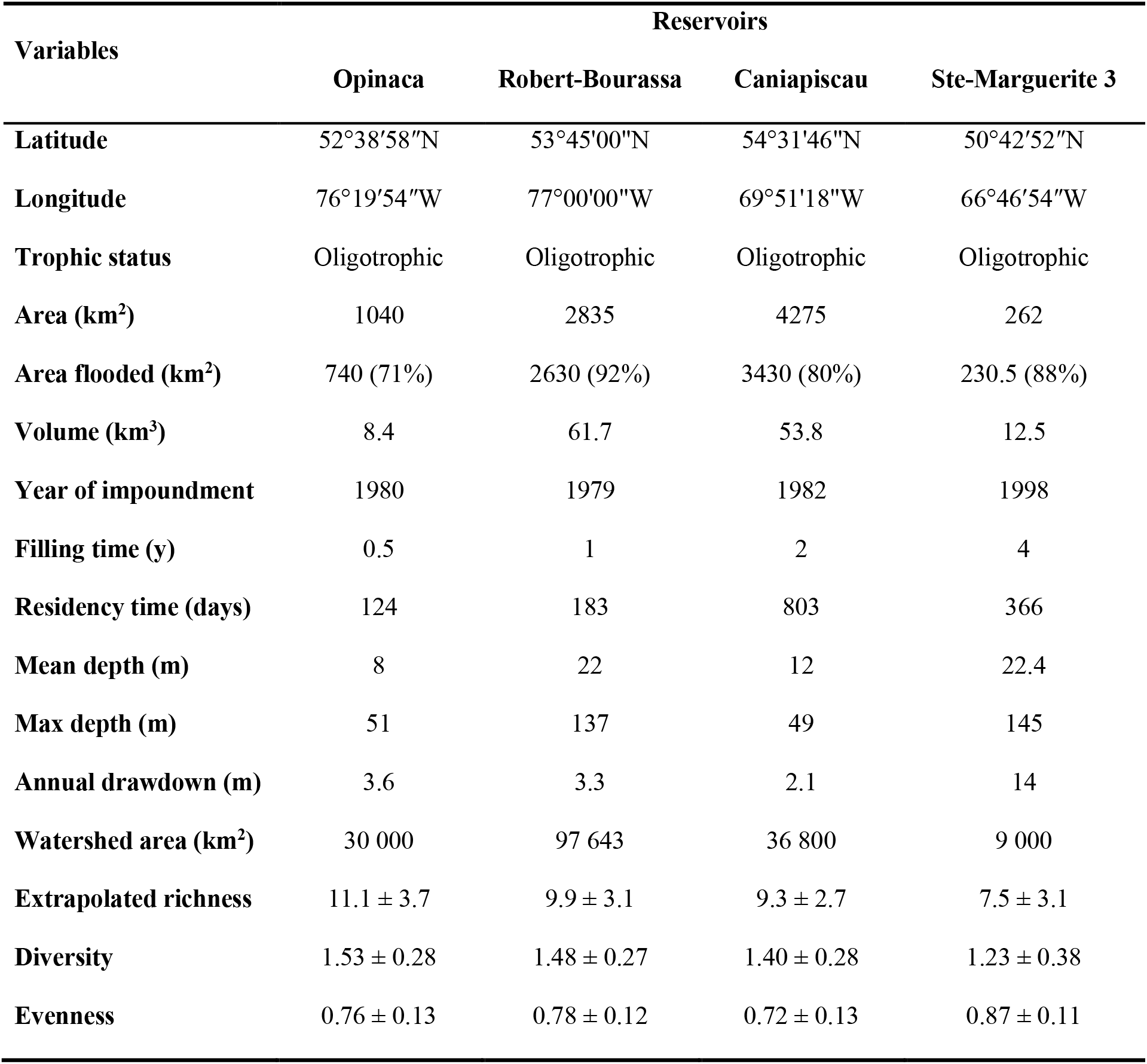
Reservoirs characteristics in the La Grande and Sainte-Marguerite 3 hydroelectric complexes. Reservoir area represents the surface covered with water at maximum pool. The area flooded represents the surface of terrestrial land flooded following impoundment and, in brackets, the percentage of the reservoir that was terrestrial land before impoundment. The extrapolated richness, diversity and evenness values are averages calculated across all upstream and downstream stations for all time points.

#### Sainte-Marguerite complex

The Sainte-Marguerite complex (SM) is located on the Moyenne-Côte Nord portion of the Canadian Shield (Eastern Québec, Canada; Fig. 1). The Sainte-Marguerite 3 reservoir (SM3) is deeper than the LG reservoirs (Table 1) and is located within a canyon shape valley. The Sainte-Marguerite river was impounded by Hydro-Québec in 1998 and took 4 years to fill. A smaller downstream reservoir (SM2) was created in 1954 by Gulf Pulp and Paper (pulp industry) and is now managed by Gulf Power Co. The Sainte-Marguerite watershed is also relatively free of anthropogenic perturbation. Fish community data were collected in two stations upstream of the SM3 dam (two UR), in one UR station in SM2 reservoir that is downstream of SM3 but cannot be classified as a “true” downstream station, and in one reference station (Table 1, Fig. 1).

### 2.2 Field sampling

In RB and OP reservoirs, nearshore fish communities were sampled annually from 1978 to 1984, and then in 1988, 1992, 1996 and 2000. In RB, the pre-impoundment period corresponds to 1978 whereas in OP this period corresponds to the years 1978 and 1979. In CA reservoir, fish communities were sampled annually from 1980 to 1982, and in the 1987, 1991, 1993, 1997 and 1999. In CA, pre-impoundment data correspond to the years 1980 and 1981. In SM3 and SM2, the fish community was sampled in 1992, 1996, 2005 and 2011, with the former two years corresponding to pre-impoundment period.

In LG complex, fish sampling occurred monthly from June to September-October, with a total of five sampling times per season in RB and OP reservoirs, and four times per season in CA until 1995. After 1995 in the LG complex, and for the whole period in SM complex, the fishing protocol was optimized to concentrate the sampling effort in July and August (Deslandes & Fortin 1994). To standardize the time series, we only used the data for the months of July and August. Four gill nets were used, set in pairs. In each pair, there was an experimental multifilament gill net (45.7 m in length x 2.4 m in depth; mesh sizes ranged between 2.5 to 10.2 cm). The second net in the pair was a gill net of uniform mesh size (either with a stretch mesh size of 7.6 cm or 10.2 cm). Each net pair was set perpendicular to shore. In one of the net pairs, the gill net with uniform mesh size was directed onshore while the other pair had the gill net with uniform mesh size directed offshore. Sampling periods lasted 48h (nets visited every 24 h) in LG complex until 1982 and 24h from 1983, and lasted 48h in the SM complex. All fish caught were counted, measured and weighed. No seine net or minnow traps were used in this sampling program and no gill nets were set in the pelagic zone. Thus, the presence and abundance of small species from the nearshore area and pelagic species were underestimated.

In LG complex, changes in water quality in the photic zone (0 - 10 m) were monitored at the same sampling stations. Water quality variables measured were average water temperature measured at every m in the photic zone, water transparency (measured as secchi disk depth), dissolved oxygen concentration, pH and specific conductivity (all measured with a Hydrolab multiprobe). Details of the methodology used in the collection and analysis of these data were presented by Fréchette (1980).

### 2.3 Statistical analyses

#### Alpha- and gamma-diversity analysis

Diversity was assessed with extrapolated species richness, Pielou's J Evenness index, and Shannon's H′ diversity index. The extrapolated species richness represents the number of species for a given standardized number of net lifts and we used the second-order jackknife index (Jack2; function specpool in the vegan R package v. 2.4-1; Oksanen *et al.* 2016). Shannon's H′ diversity index takes evenness and species richness into account and quantifies the uncertainty in predicting the species identity of an individual that is taken at random from the dataset. Pielou's J′ Evenness index ranges from near 0 (indicating pronounced dominance) to near 1 (indicating an almost equal abundance of all species).

To examine changes in diversity metrics over time in impacted stations in relation to reference sites, we used General Linear Mixed Effects Models (glmm; applying the lme function from the nlme package v. 3.1-128). Here, we were interested to compare the slopes (*i.e.*, interaction term between time since impoundment [TSI] and impacted vs. reference sites [RI], Table 2). We examined the effect of river impoundment on diversity metrics at three spatial scales: at the hydroelectric complex scale (gamma-diversity; pooling data for all impacted stations in each complex), at the reservoir scale (gamma-diversity; pooling data from impacted sampling stations in each reservoir) and at the sampling station scale (alpha-diversity). To control for spatio-temporal dependence, we used random factors where sampling stations were nested within reservoirs: ~1 + TSI |STATION/RES (where RES stands for reservoir identity). We also used an autoregressive correlation structure (corAR1) to control for temporal autocorrelation. We determined the autoregressive process in each time series by plotting each time series and by observing the autocorrelation function (ACF) and the partial autocorrelation function (PACF) on detrended data using an autoregressive integrated moving average model diagnostic (astsa package v. 1.3 in R; Stoffer 2014). Errors followed an autoregressive process of degree 1. Our glmms with the autocorrelated structure did not perform better than those without based on AICc scores (Burnham & Anderson 2002). We present only the glmms without the autocorrelated structure.

**Tabel 2.**
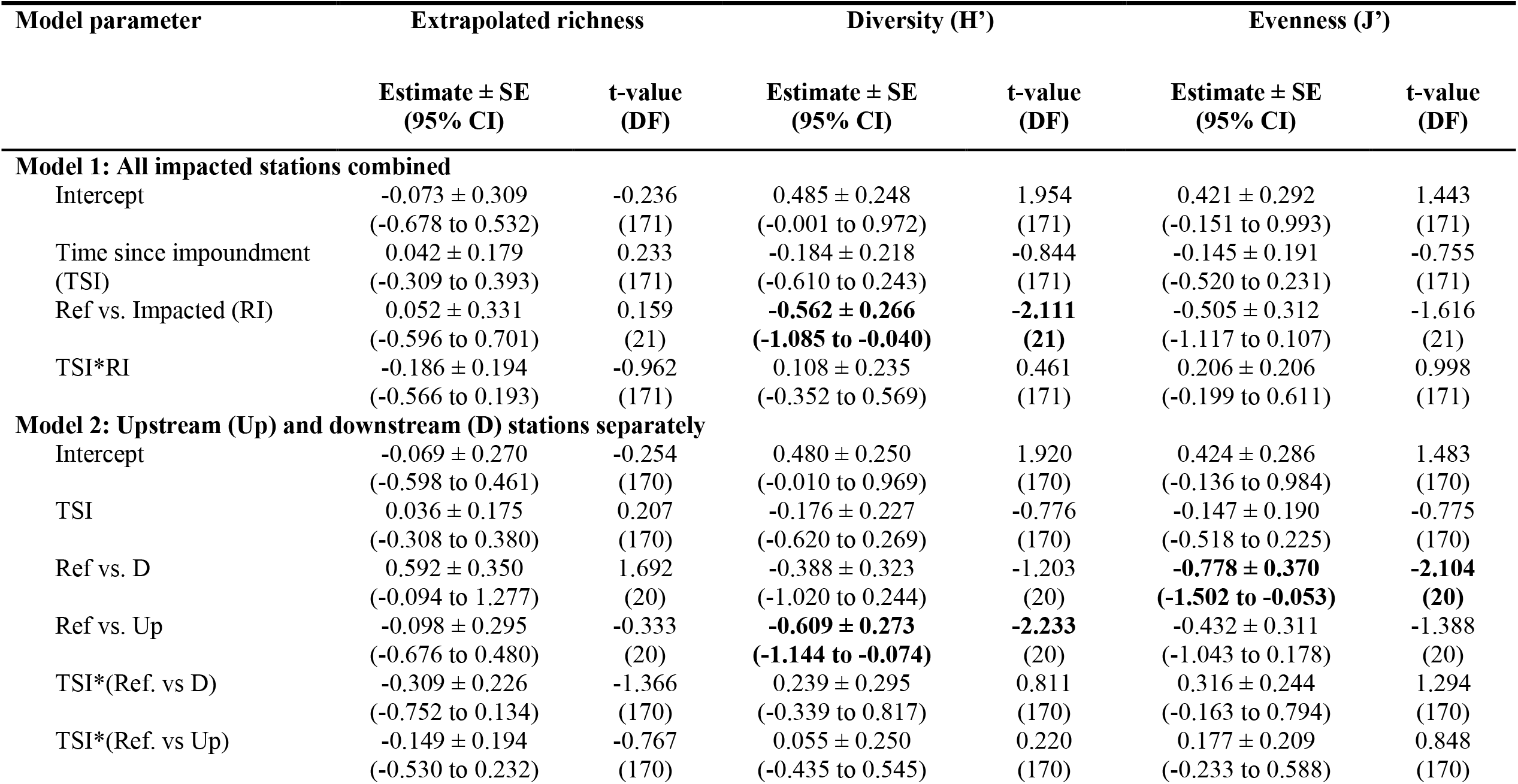
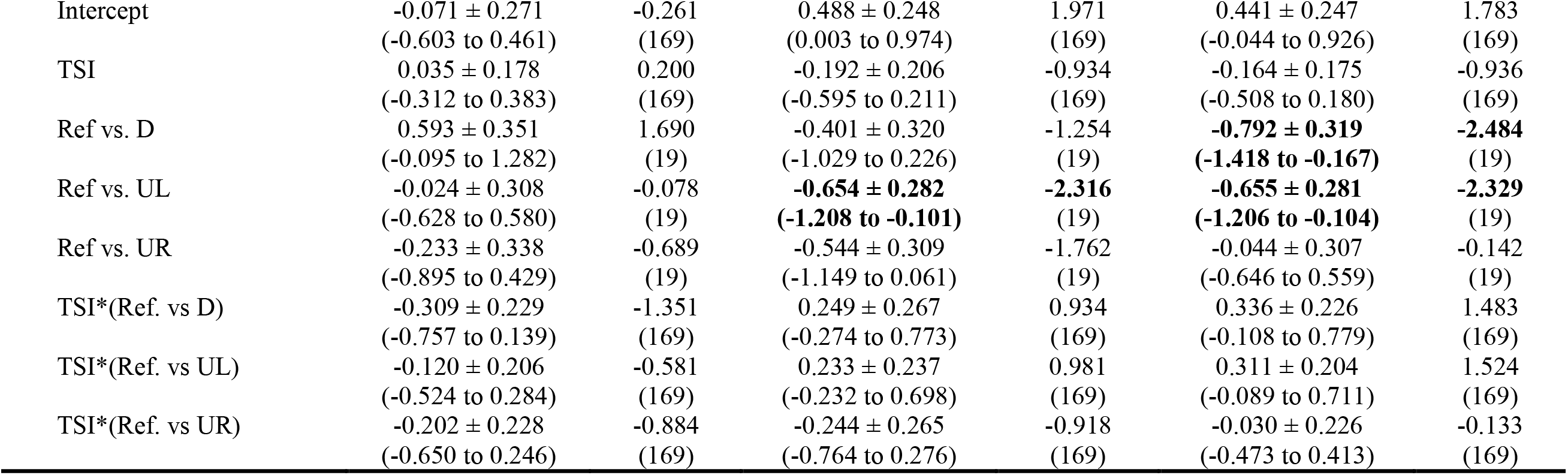
Estimate ± Standard error (SE), 95% Confidence intervals, t-values and degrees of freedom (DF) of model parameters used to predict change in extrapolated richness, diversity and evenness in La Grande hydroelectric complex. General linear mixed effects models were used to evaluate the effect of time since impoundment, stations categories (Impacted stations vs. reference sites) and their interaction on diversity metrics. Predictors not including 0 within their 95% CI are in bold. Reference sites are used as contrasts in the models.

#### Beta-diversity analysis

To test species turnover rate over time and space, we computed Local Contributions to Beta-Diversity (LCBD; using the *beta.div* function in R available at <u>http://adn.biol.umontreal.ca/~numericalecology/FonctionsR/</u>) and Species Contributions to Beta-Diversity (SCBD) indices on Hellinger-transformed species abundance community matrices (Legendre and De Cáceres 2013). LCBD values indicate how unique is any fish composition of a site relative to other comparable sites by assessing its contribution to the total variation in fish composition in space and/or time. SCBD indicates how large of a contribution is a species has to overall beta diversity in the dataset (Legendre & De Cáceres 2013; Legendre & Gauthier 2014). For details about the calculation of LCBD and SCBD and the. We computed LCBD at the complex scale, at the reservoir scale, and at the sampling station scale and SCBD at the sampling station scale. At the sampling station scale, each station was evaluated separately, so the turnover rate was in relation to time only.

### 2.4 Variation partitioning

We examined fish species assemblages at the three spatial scales using unbiased variation partitioning based on RDAs (Redundancy Analysis) and adjusted R^2^ (Peres-Neto *et al.* 2006) with the *varpart* function in the vegan package (v. 2.4-1). We used the forward selection procedure of Blanchet, Legendre & Borcard (2008). With variation partitioning analyses, the overall variation in species assemblages can be divided into “fractions” attributable to different data matrices as well as combinations of these matrices (*i.e.*, shared variation). Here, we used four matrices: time since impoundment [TSI; *i.e.,* including years before and after impoundment], spatial heterogeneity [SH; latitude, longitude and identity of each sampling station and reservoir], water quality variables [WQV; water transparency, dissolved oxygen, pH, conductivity and temperature] and fishing gear [G]. The total variation of species assemblages was decomposed into 15 fractions at the complex and reservoir scales, and eight fractions at the sampling station scale because the [SH] matrix is irrelevant at the sampling station scale. We used the Hellinger-transformed abundance values of species. To produce the most parsimonious model in RDAs, we performed forward selection using the double stopping criteria (*ordiR2step* function in the vegan R package v. 2.4-1; Blanchet, Legendre & Borcard 2008). Because of a small sample size, these analyses were not possible in the SM complex (only 3 to 5 observations per sampling station).

## 3. Results

### 3.1 Changes in alpha-, beta- and gamma-diversity

Overall, OP reservoir had a higher mean extrapolated richness and diversity, and SM3 had a lower richness and diversity than RB and CA reservoirs (Table 1). Downstream stations generally had higher extrapolated richness than upstream stations, but did not differ in diversity and evenness (Fig. 2). Across all scales and categories of impacted stations (U vs D and UL, UR vs. D), the temporal trends in richness, diversity and evenness in impacted stations were weak and comparable to those observed in reference sites for both complexes (complex scale; Fig. 2, Table 2, reservoir scale; Table S1, sampling station scale; Tables S2-S5).

**Figure 2.**
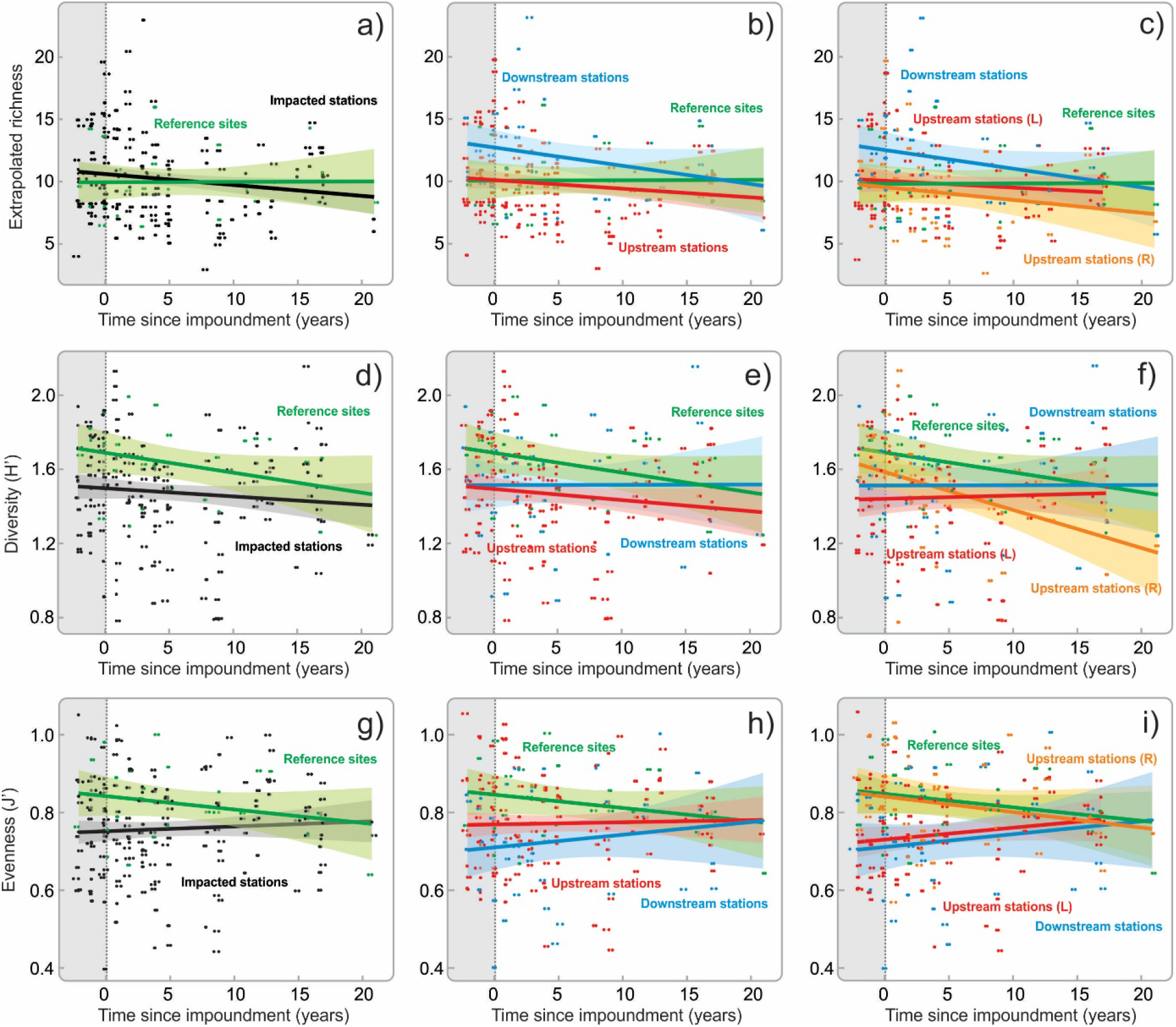
Variation in extrapolated richness, diversity (H’) and evenness (J’) over time in impacted and reference stations at the LG complex level. Changes in diversity metrics over time in references sites (green) were compared with impacted stations (all categories combined from all reservoirs) in a, d, g panels, with impacted stations upstream and downstream of the dams in b, e, h panels, and with upstream stations that were lakes before being a reservoir (UL) and those that were a river before (UR) in c, f, i panels.

For completeness, we also examined the temporal trends in impacted stations only, without comparison with reference sites. At the complex scale, we did not detect any temporal trend in diversity metrics when categories of impacted stations were combined (Model 1; Table S6). When station categories were added in the model (*i.e.*, U vs. D, or UR, UL vs. D), richness marginally decreased over time (Models 2 and 3; Table S6). This trend was strongly driven by the low richness values observed in 2000 in RB (lower fishing effort in this one year). When this data point was excluded from the analysis, the trend was not significant anymore. At the reservoir scale, we found some decreasing temporal trends in RB (Table S7, Models 1, 2, and 3) but found no temporal trends in the other reservoirs (Table S7).

The lack of strong temporal trends in alpha- and gamma-diversity was echoed by an absence of clear beta-diversity patterns across space and time at either the complex (Fig. 3 a and Fig. S1) and reservoirs scale (and Fig. S2). At both scales, relatively few Local Contribution to Beta-Diversity (LCBD) values were significant, and the weight of LCBD values did not relate to impoundment, nor to the impacted stations. However, when beta-diversity analyses were conducted at the sampling station scale (*i.e.,* only comparing any one site to itself through time), many of the significant LCBD values were apparent in upstream stations during and shortly after filling (Fig. 3 b), showing a higher species turnover rate during this period.

**Figure 3.**
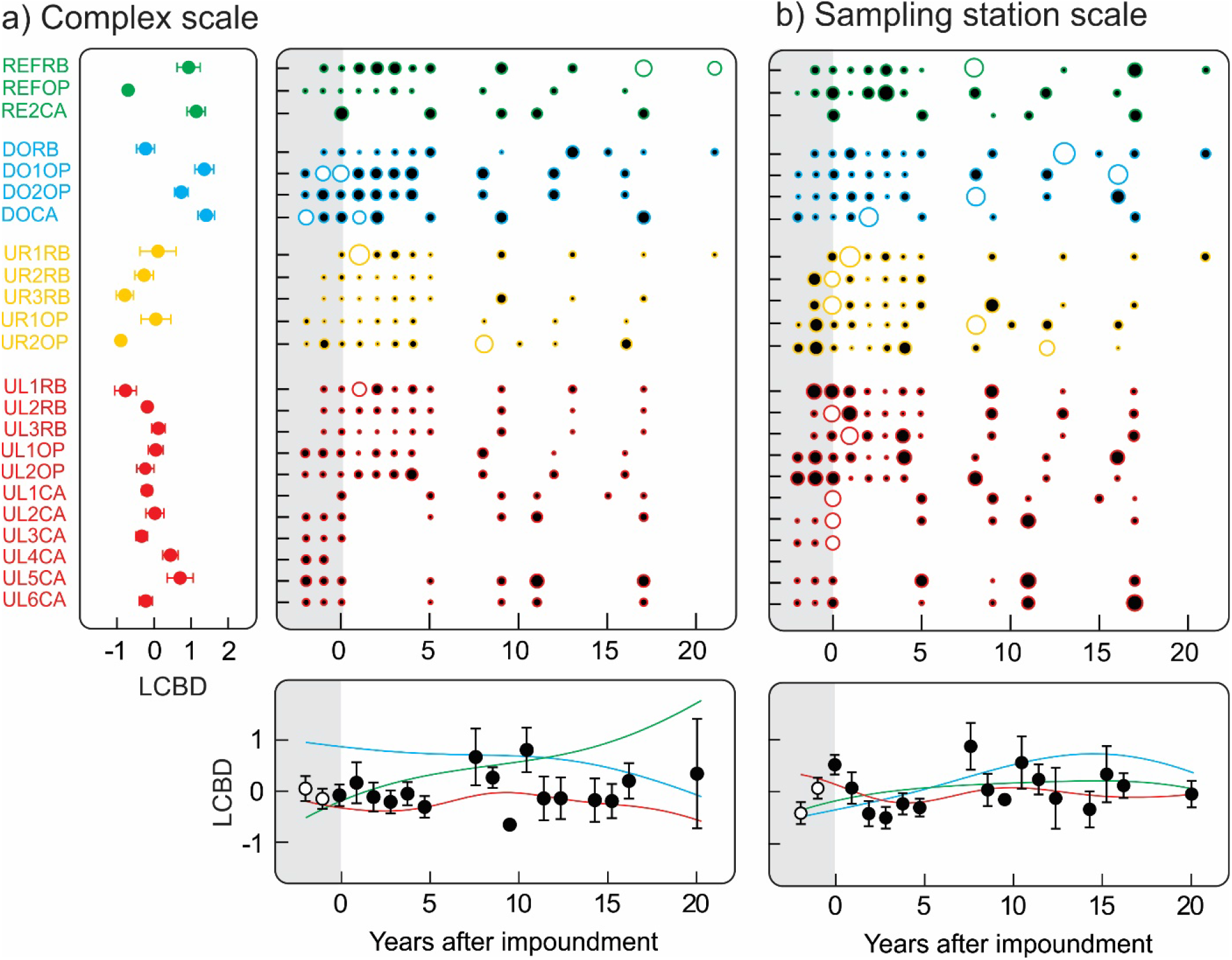
Local contribution to beta-diversity (LCBD) per station and year at: a) the LG complex level and b) the sampling station level. Circle areas are proportional to the LCBD values. Circles filled in white indicate significant LCBD at p<0.05. The lower panels represent mean values of LCBD per year with a distance weights least square (DWLS) curve fit. The right panel represents mean values of LCBD per station for the analysis at the complex level. Stations with a label starting with “UL” represent stations that were lakes before being a reservoir and those with a label starting with “UR” were rivers or stream before being a reservoir. Reference sites are in green, downstream stations in blue and upstream stations in yellow UR) and red (UL).

### 3.2 Drivers of the shift in species assemblage

At the complex and reservoirs scales, spatial heterogeneity (SH) among sampling stations was the main driver structuring fish assemblages (Fig. 4 a-b, Table S8). The effect of impoundment only became a dominant predictor at the sampling station scale (Fig. 4 c, Table S9). At the scale of the LG complex, SH explained 45% of the variation across all shared fractions (25% explained by SH alone) and a significant proportion of the variation was shared with water quality variables (WQV; 15%; Fig. 4 a). A similar pattern was observed at the reservoir scale (Fig. 4 b; Tables S8 and S9). At the scale of the sampling stations, most of the variation was explained by the shared effect of TSI and WQV (24%; Fig. 4 c and Table S3), which suggests that fish responded very locally to impoundment and in a large extent to changes in water quality associated with impoundment.

**Figure 4.**
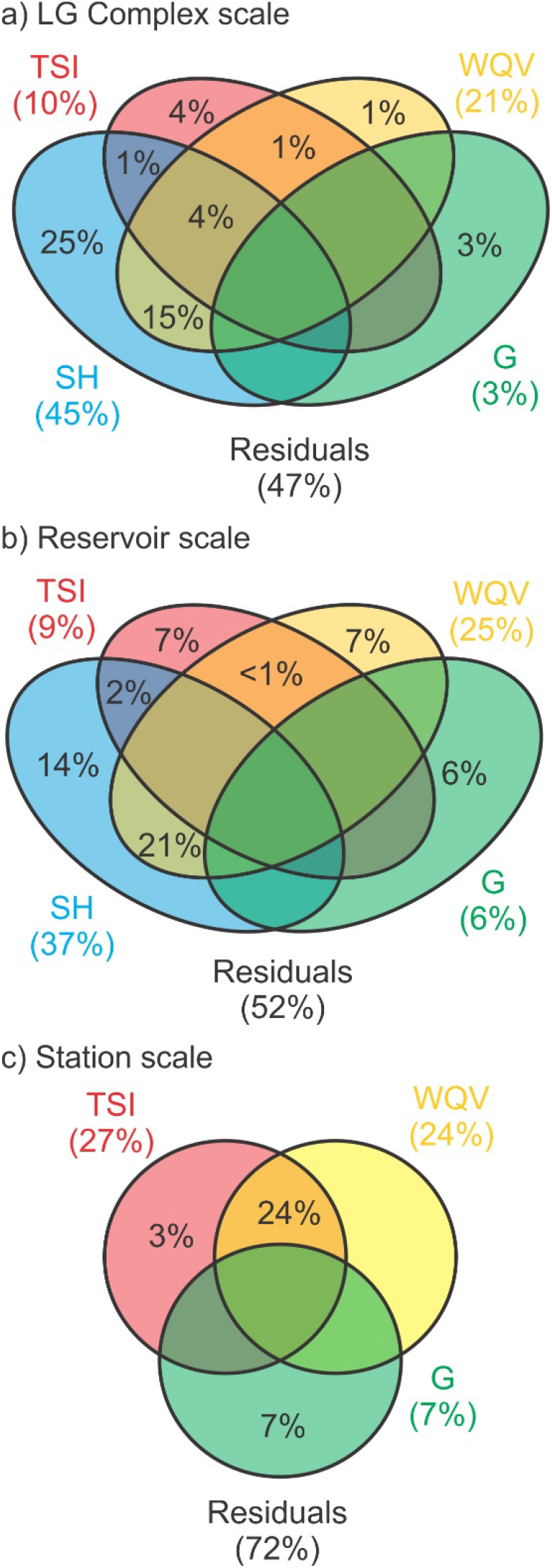
a) Variation partitioning analysis showing the contribution of four matrices (Time since impoundment [TSI], Spatial heterogeneity [SH], Water quality variables [WQ] and fishing gear [G]) to explain the variation in fish species assemblages at the a) LG complex level, b) at the reservoir level (average across reservoirs, see Table S7 for the breakdown per reservoir), and c) at the sampling stations (average across sampling stations, see Table S8 for the breakdown). All analyses included reference sites.

### 3.3 Species affected by impoundment

The effect of impoundment on species differed among reservoirs, and among the categories of sampling stations (Fig. 5 and Fig. 6). In several upstream stations, we observed a shift from a catastomids-dominated community (longnose sucker, *Catostomus catostomus* and white sucker, *C. commersonii*) toward a pike-coregonids (northern pike, *Esox lucius*, whitefish *Coregonus clupeaformis* and cisco, *C. artedi*) community after impoundment (Fig. 6). This shift was supported by high contribution to beta-diversity (SCBD) for these species in upstream stations (Fig. 5 b, c, d and e). Changes in species assemblages in upstream stations appear to have mostly occurred within the first 5 years of impoundment (Fig. 6). In downstream stations, no consistent pattern was observed but the marked changes were a decrease of the lake sturgeon (*Acipenser fulvescens*) and an increase in walleye (*Sander vitreus*) in OP (Fig. 6 f), and a decrease in abundance of the brook trout (*Salvelinus fontinalis*) in CA. These patterns were echoed by the SCBD values (Fig. 5 d, d and e). Reference sites were more stable but also experienced some changes in fish community structure, with a fluctuating dominance between two predators in RB (*i.e.,* walleye and burbot, *Lota lota;* Fig. 6 g), and between the lake trout (*Salvelinus namaycush*) and the two catastomids in CA (Fig. 6 i).

**Figure 5.**
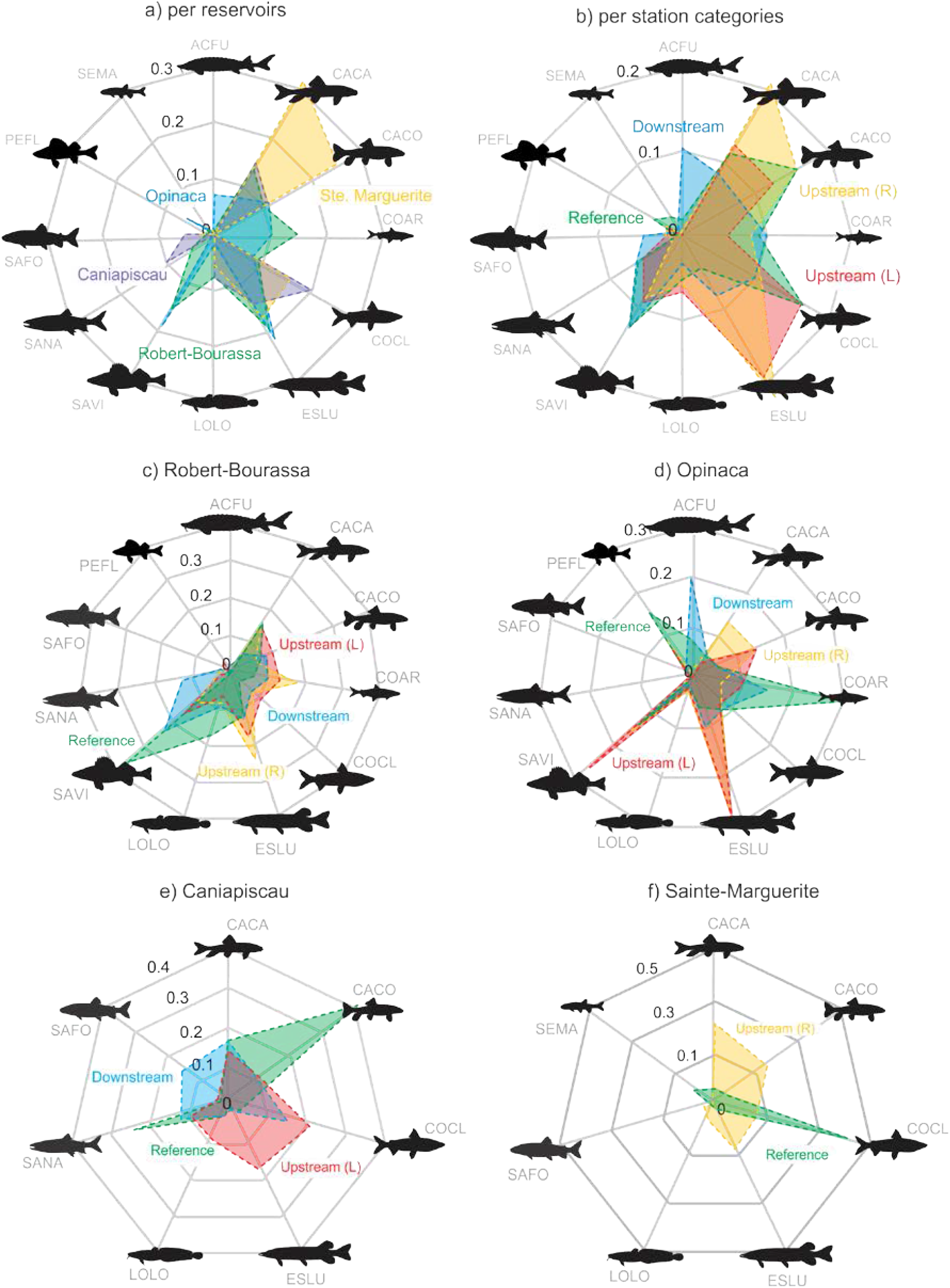
Radar charts of species contributions to beta-diversity (SCBD) computed for each sampling stations, and pooled: a) per reservoirs, b) per categories of sampling stations, for c) Robert-Bourassa, d) Opinaca and e) Caniapiscau and f) Sainte-Marguerite. Only the ten most common species are pictured.

**Figure 6.**
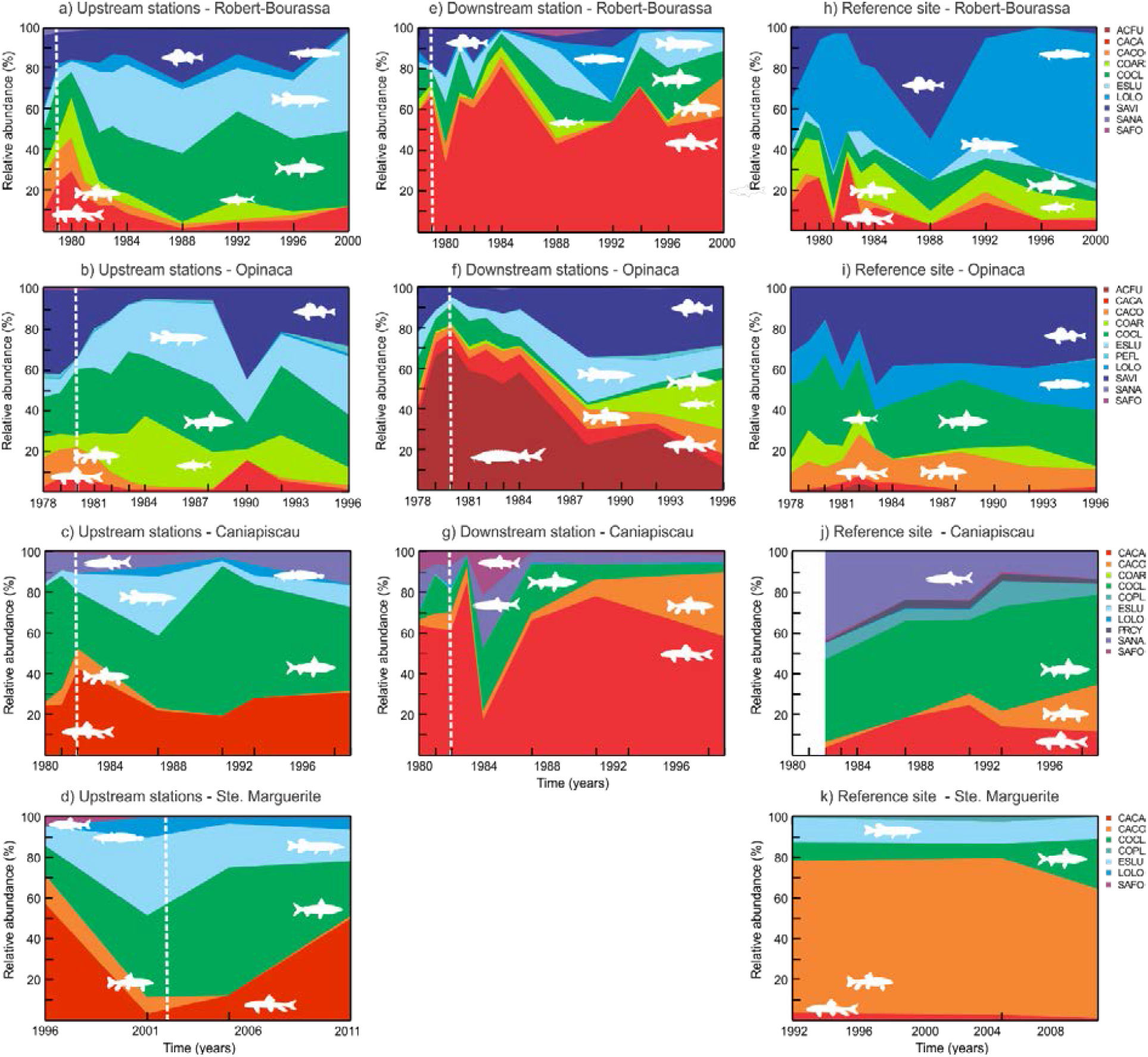
Changes in relative abundance over time of the most common species (>5% of total catch) in sampling stations of LG reservoirs and SM3 reservoir. Relative abundance in upstream stations for each reservoir are in a, b, c and d panels, in downstream stations in e, f and g panels and in reference sites in h, i, j k panels. The white dashed line represents the start of reservoir filling. See Table S9 for the name of the species related to the four letters code.

## 4. Discussion

### 4.1 Fish community response to impoundment

There is an extensive literature on fish community responses to impoundment in temperate and tropical reservoirs, but little was known about boreal reservoirs (but see Tereshchenko & Strel’nikov 1997; Sutela & Vehanen 2008). Our analyses showed that in four remote large boreal reservoirs, there were no significant temporal trends in fish alpha- and gamma-diversity at three spatial scales. No native species were lost, and non-native fish did not colonise our boreal study systems. Our work is in marked contrast to the tropic, where there is evidence of a net loss of species ( Liew, Tan & Yeo 2016). As such, there appears to be an important heterogeneity in fish diversity responses to impoundment. However, almost all studies to date (including ours) have shown a general change in fish assemblage after impoundment.

From the literature, four main mechanisms have been suggested to cause a change in fish assemblage in reservoirs: 1) shift from a lotic to a lentic environment upstream of the dam, 2) dams as barriers to free movement, 3) alteration of the natural hydrological regime, and 4) higher susceptibility of reservoirs to be invaded by non-native species. Are these mechanisms comparable across latitudes? The shift from lotic to lentic conditions upstream of the dam represents an extreme transformation to fish habitats and can exert a suite of selective pressures not experienced by fish during their evolutionary history. This is especially true in the tropics where fish have evolved in flowing waters (Gomes & Miranda 2001) and may lack the morphological and behavioral characteristics, or the reproductive strategy and plasticity to successfully occupy the new lentic habitats (Gomes & Miranda 2001; Agostinho, Pelicice & Gomes 2008). Given the predominance of large rivers and streams in the tropics and temperate environments, significant losses in richness in these regions have been attributed to the transformation of ecosystems in lentic ones (Martinez *et al.* 1994; de Mérona, Vigouroux & Tejerina-Garro 2005; Sá-Oliveira *et al.* 2015; Lima *et al.* 2016). In boreal regions, both large lakes and rivers are common (Messager *et al.* 2016) and the evolutionary young fish species found in this region appear to be somewhat resilient to river impoundment. The creation of new lentic habitats upstream of the dam, captured by the TSI variable in our study, is the most plausible driver of the shift in assemblages, but did not wipe out any species.

Dams can also block migratory route of diadromous species and alter seasonal migration of potamodromous species. Local losses or reduction in abundance of migratory species has been attributed to river fragmentation by dams in tropical and temperate regions (Reyes-Gavilán *et al.* 1996; Galat *et al.* 1998; Gehrke, Gilligan & Barwick 2002; Okada, Agostinho & Gomes 2005; Sá-Oliveira *et al.* 2015; Pelicice, Pompeu & Agostinho 2015; Lima *et al.* 2016). In our boreal systems, dams did not appear to be a major barrier to migration or movement for most species because focal fishes were not diadromous and do not undertake long spawning migration (Table S10). Moreover, the barrier effect might also have been minimized as most dams were built on pre-existing obstacles that were already impassable for fish (*i.e.,* high waterfall). However, studies of this nature should be pursued in the future because of the high occurrence of anadromy in some boreal regions (McDowall 2008).

The effects associated with altering the natural hydrological regimes on fish communities depend in part on reservoir morphometry and on the magnitude of the alteration that is related to reservoir management (*e.g.*, magnitude of drawdown, discharge and hypo- vs. epilimnitic water release). As noted in previous works, the magnitude of change in discharge and drawdown can have divergent effects. For example, a 76% decrease in discharge in the Canadian River strongly affected fish assemblages downstream of the Ute dam (Ute reservoir, New Mexico, USA), but a 36% decrease in discharge did not have significant effects downstream of Sanford dam along the same river (Lake Meredith reservoir, Texas, USA; Bonner & Wilde 2000). In our boreal ecosystems, despite the diversion of some rivers (decrease in discharge of up to 90%), only the lake sturgeon in Opinaca appear to be really strongly affected (Figs 5 and 6).

Intentional (*e.g.*, fishing bait) or unintentional introduction (*e.g.*, flooding creates new connection between water bodies) of non-native species in reservoirs can promote a shift away from native-dominated fish communities (Rodriguez Ruiz 1998; Gido *et al.* 2002; Johnson, Olden & Vander Zanden 2008; Clavero & Hermoso 2010). As an extreme example, the introduction of a voracious non-native predator (the peacock-bass; *Cichla kelberi*) in Rosana reservoir (Paraná river basin), decreased fish richness by 80% after only three years (Pelicice and Agostinho 2008). River basins where endemic species are abundant might be particularly vulnerable (Dudgeon *et al.* 2006). In our boreal reservoirs, no non-native species has been observed, and no endemic species were present in either the LG or SM complexes. The remote location of our focal reservoirs also likely contributed to the lack of establishment of non-native fishes.

The time it takes for fish communities to stabilize after impoundment is highly variable among studies. It has been reported to be either quick (*i.e.,* within five years; Martinez *et al.* 1994), or much longer (more than 10 years; Quinn & Kwak 2003; Říha *et al.* 2009), highlighting the need for data span decades after impoundment. Some states or phases can also be transient. Říha *et al.* (2009) documented a five-phase succession in fish species with European reservoirs aging. The time needed for the fish assemblages to stabilize will depend on fish behavior, life history trait and adaptability, the stability of the food web, the strength of trophic interactions, and on the management and operation of the dam and reservoir. If the dominant mechanisms are related to reproduction and recruitment through the strength of year classes, the effect may take years to be detectable. Species with some specific life history traits (*e.g.*, late age at first reproduction), or positioned at higher trophic levels may have delayed responses to impoundment. If the dominant mechanisms are through movement and redistribution due to river fragmentation and change in habitat quality, then shifts can be detected quickly.

### 4.2 Multi-scale approach and study design

Equipped with fish assemblage data collected over decades after impoundment, and across a large spatial network of sites in a remote boreal region, this study is unique in providing the most data-rich analysis to date, and in its ability to isolate the effect of impoundment from other factors that co-occur with hydroelectricity projects. Great insights are achieved when multiple scales are considered because patterns observed in communities at a given scale are often the consequence of a complex interplay between various processes occurring at multiple scales (Wiens 1989; Whittaker, Willis & Field 2001). In this study, changes in fish assemblages in response to impoundment were only detectable at the sampling station scale (Figs. 3 and 4). At the complex and reservoirs scales, fish assemblage shifts were largely masked by some other larger scale ecological processes (*i.e.,* diverse habitat types and natural barriers to movement leading to different fish communities), which highlights the importance of a multi-scale approach to evaluate the potential of anthropogenic impacts on aquatic ecosystems.

Scale matters, but having different categories of stations, reference sites, and time series covering the periods before and after impoundment are equally important considerations to understand the effects of impoundment on fish communities. We found the strongest shifts in species assemblages in upstream stations (Fig 6), relative to references and downstream stations, which clearly points to the impact of impoundment vs. regional environmental change. Finally, time series should cover the period before and after impoundment, and preferably time series should be long enough to cover the non-equilibrium trophic surge and reach the new ecosystem equilibrium (Grimard & Jones 1982; Turgeon *et al.* 2016).

## 5. Conclusions

By using a network of sites with minimal confounding factors, and by conducting our analyses at three spatial scales, we have provided strong empirical evidence that impounding large rivers in these boreal ecosystems did not affect diversity, but resulted in a clear shift in fish assemblages. Changes in fish assemblages to impoundment were most clearly detected with our ordination and beta-diversity analyses conducted at the scale of the sampling station. Given the strength of our multi-scale approach in providing a complete perspective on the scale at which river impoundment affect fish community, we caution against large scale extrapolations and correlation studies that may underestimate or mask anthropogenic effects on aquatic ecosystems. Reservoirs are now dominant features of many landscapes, and they will become even more common in the coming years, especially in tropical regions (Zarfl *et al.* 2014; Winemiller *et al.* 2016). Identifying which mechanisms related to impoundment and river regulation affect species, evaluating the strength of their effects, and how they vary across regions can assist in implementing mitigation measures and in minimizing biodiversity loss.

## Authors’ Contributions

KT, CT and IGE conceived the idea, KT analysed the data; KT led the writing of the manuscript. All authors contributed critically to the drafts and gave final approval for publication.

---

**Tabel S1.**
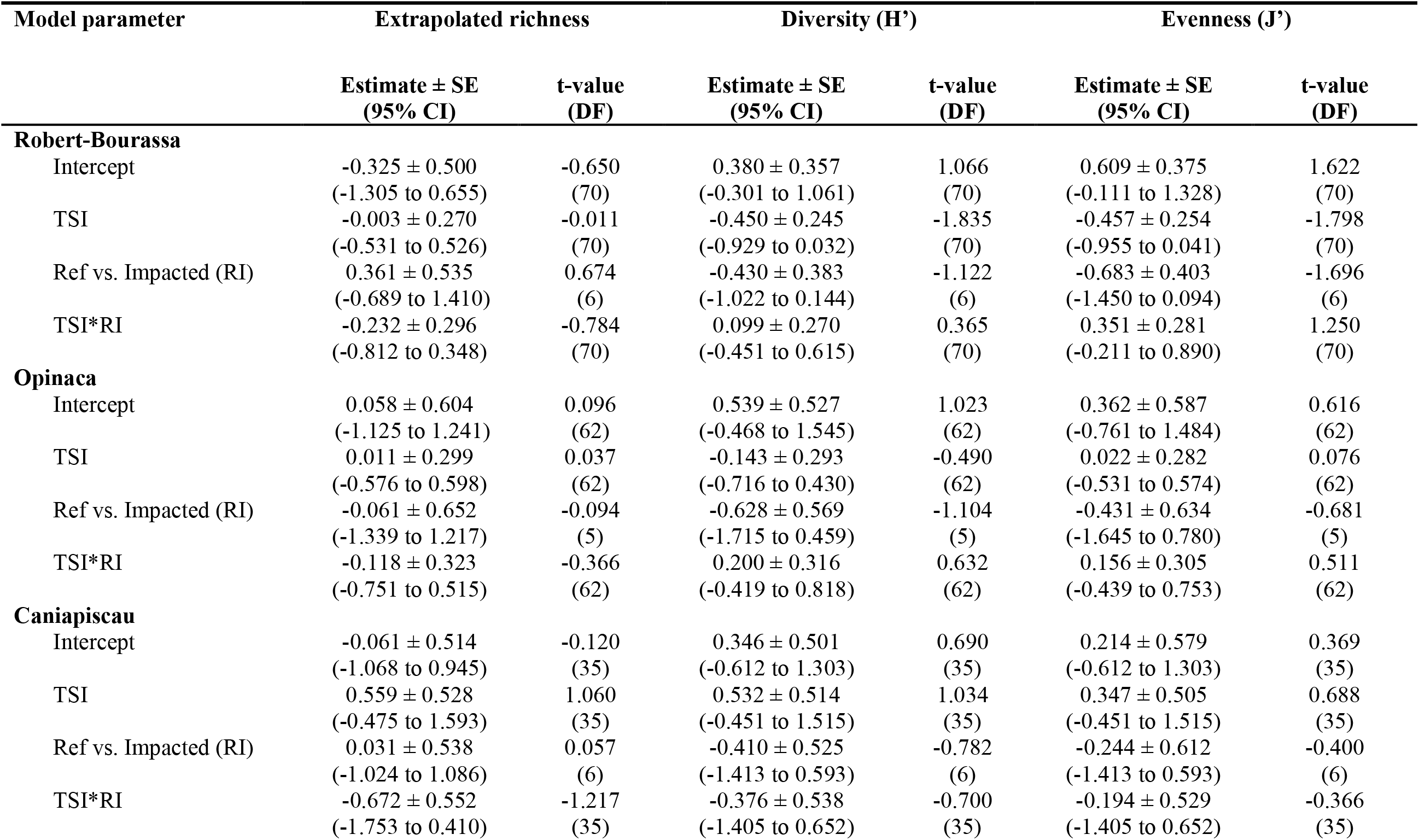
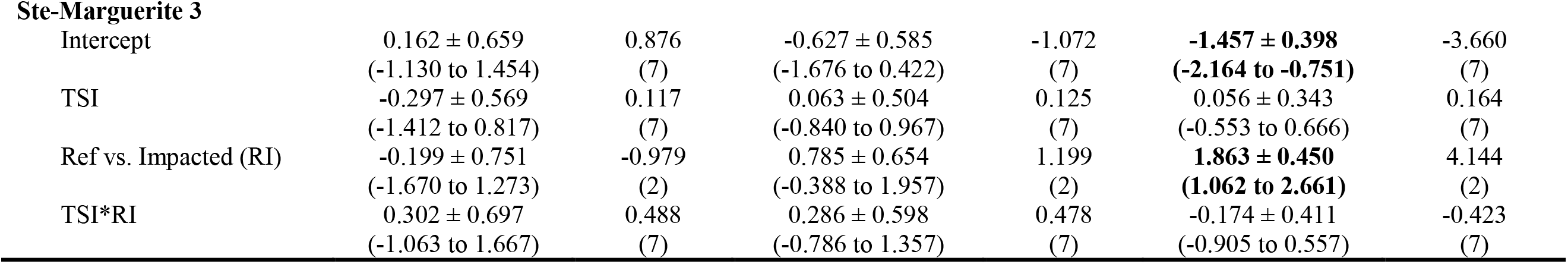
Analysis at the reservoir scale. Estimate ± Standard error (SE), 95% Confidence intervals and t-values, and degrees of freedom (DF) of model parameters used to predict change in extrapolated richness, diversity and evenness in La Grande complex reservoirs and Ste-Marguerite 3 reservoir. Generalized mixed effects models were used to evaluate the effect of time since impoundment, stations categories and their interaction on diversity metrics. Reference sites are used as contrasts.

**Tabel S2.**
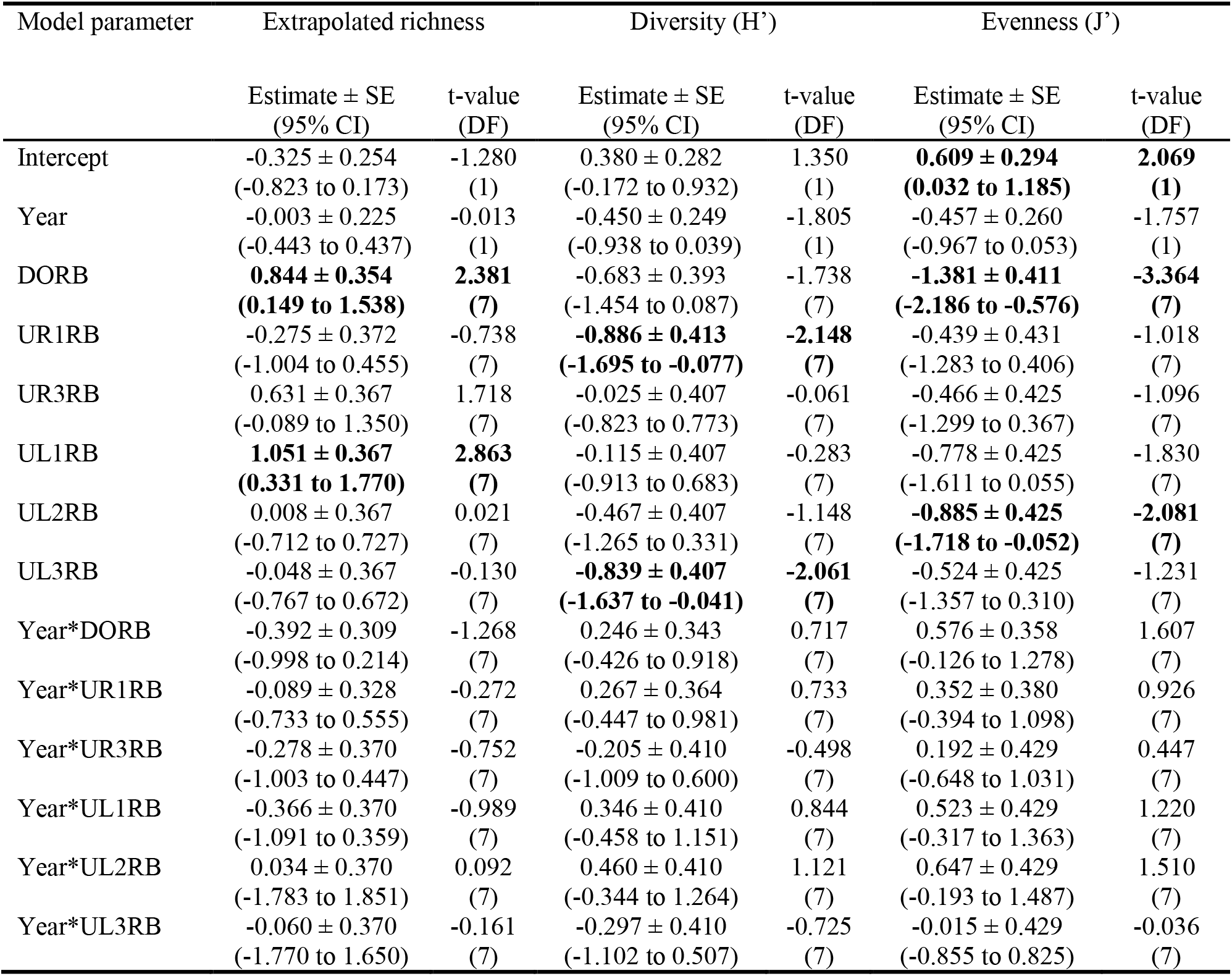
Sampling stations in Robert-Bourassa. Estimate ± Standard error (SE), 95% Confidence intervals (95% CI) and t-values of model parameters used to predict change in extrapolated richness, diversity (Shannon’s H’) and evenness (Pielou J’) in impacted stations in Robert-Bourassa reservoir when compared to reference site. General linear models were used to evaluate the effect of time, stations and their interaction on diversity metrics. The reference site is used as a contrast in the model.

**Tabel S3.**
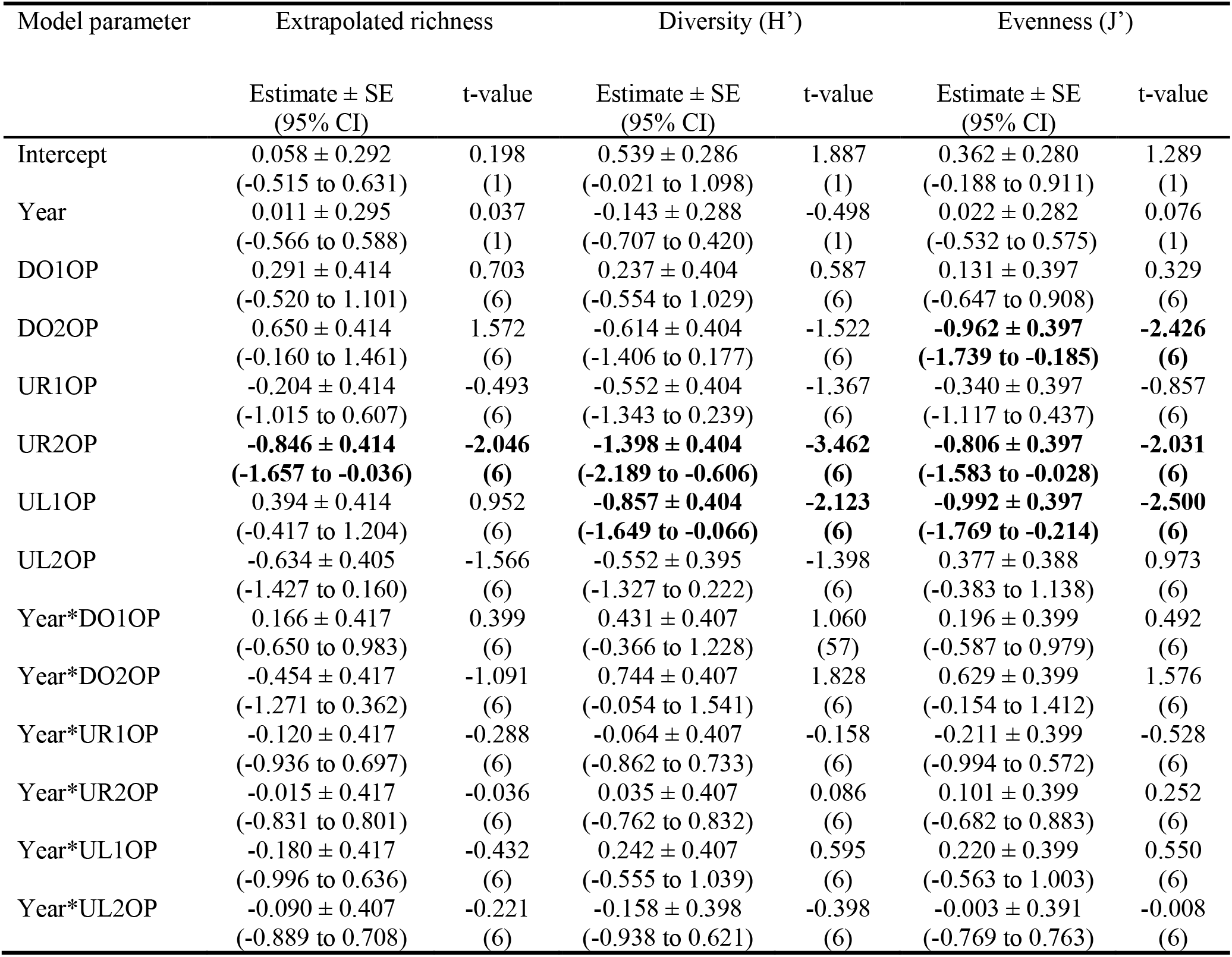
Sampling stations in Opinaca. Estimate ± Standard error (SE), 95% Confidence intervals (95% CI) and t-values of model parameters used to predict change in extrapolated richness, diversity (Shannon’s H’) and evenness (Pielou J’) in impacted stations in Opinaca reservoir when compared to reference site. General linear models were used to evaluate the effect of time, stations and their interaction on diversity metrics. The reference site is used as a contrast in the model.

**Tabel S4.**
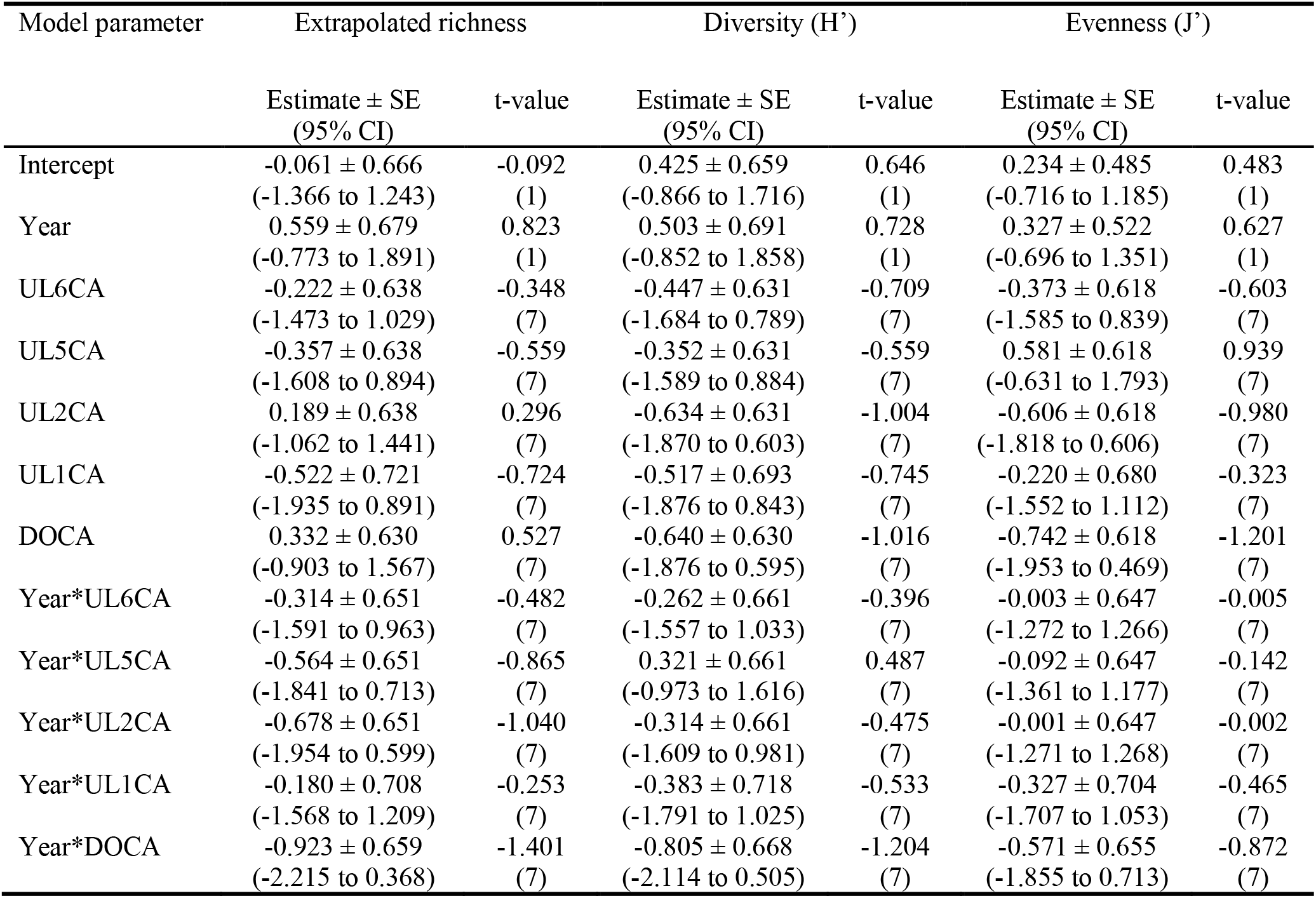
Sampling stations in Caniapiscau. Estimate ± Standard error (SE), 95% Confidence intervals (95% CI) and t-values of model parameters used to predict change in extrapolated richness, diversity (Shannon’s H’) and evenness (Pielou J’) in impacted stations in Caniapiscau reservoir when compared to reference site. General linear models were used to evaluate the effect of time, stations and their interaction on diversity metrics. The reference site is used as a contrast in the model.

**Tabel S5.** Sampling stations in Sainte-Marguerite. Estimate ± Standard error (SE), 95% Confidence intervals (95% CI) and t-values of model parameters used to predict change in extrapolated richness, diversity (Shannon’s H’) and evenness (Pielou J’) in impacted stations in Sainte-Marguerite reservoir when compared to reference site. Generalized mixed effects models were used to evaluate the effect of time, stations and their interaction on diversity metrics. The reference site is used as a contrast in the model.

| Model parameter | Extrapolated richness | | Diversity ($H'$) | | Evenness ($J'$) | |
| --- | --- | --- | --- | --- | --- | --- |
| | Estimate ± SE<br>(95% CI) | t-value | Estimate ± SE<br>(95% CI) | t-value | Estimate ± SE<br>(95% CI) | t-value |
| Intercept | 0.162 ± 0.843<br>(-0.215 to 0.538) | 0.192<br>(1) | -0.627 ± 0.696<br>(-1.991 to 0.737) | -0.901<br>(1) | -1.457 ± 0.425<br>(-2.289 to -0.625) | -3.433<br>(1) |
| Year | -0.297 ± 0.727<br>(0.504 to -1.099) | -0.409<br>(1) | 0.063 ± 0.600<br>(-1.112 to 1.238) | 0.105<br>(1) | 0.056 ± 0.366<br>(-0.660 to 0.773) | 0.154<br>(1) |
| UR3SM | -0.443 ± 1.661<br>(0.081 to -0.966) | -0.267<br>(3) | 0.746 ± 0.888<br>(-0.994 to 2.486) | 0.840<br>(3) | 1.767 ± 0.562<br>(0.666 to 2.868) | 3.145<br>(3) |
| UR2SM | -0.224 ± 1.226<br>(0.135 to -0.582) | -0.183<br>(3) | 0.698 ± 0.997<br>(-1.256 to 2.652) | 0.700<br>(3) | 1.565 ± 0.608<br>(0.373 to 2.757) | 2.574<br>(3) |
| UR1SM | 0.155 ± 1.129<br>(-0.114 to 0.423) | 0.137<br>(3) | 0.998 ± 0.923<br>(-0.812 to 2.807) | 1.081<br>(3) | 2.177 ± 0.563<br>(1.073 to 3.281) | 3.864<br>(3) |
| Year* UR3SM | 0.417 ± 1.726<br>(-0.057 to 0.891) | 0.242<br>(3) | 0.522 ± 0.824<br>(-1.092 to 2.136) | 0.634<br>(3) | -0.080 ± 0.516<br>(-1.091 to 0.931) | -0.155<br>(3) |
| Year* UR2SM | 0.320 ± 1.184<br>(-0.209 to 0.849) | 0.270<br>(3) | 0.355 ± 0.977<br>(-1.560 to 2.269) | 0.363<br>(3) | 0.207 ± 0.596<br>(-0.961 to 1.375) | 0.348<br>(3) |
| Year* UR1SM | -0.122 ± 1.163<br>(0.084 to -0.328) | -0.105<br>(3) | -0.237 ± 0.960<br>(-2.118 to 1.644) | -0.247<br>(3) | -0.677 ± 0.585<br>(-1.824 to 0.471) | -1.156<br>(3) |

**Tabel S6.**
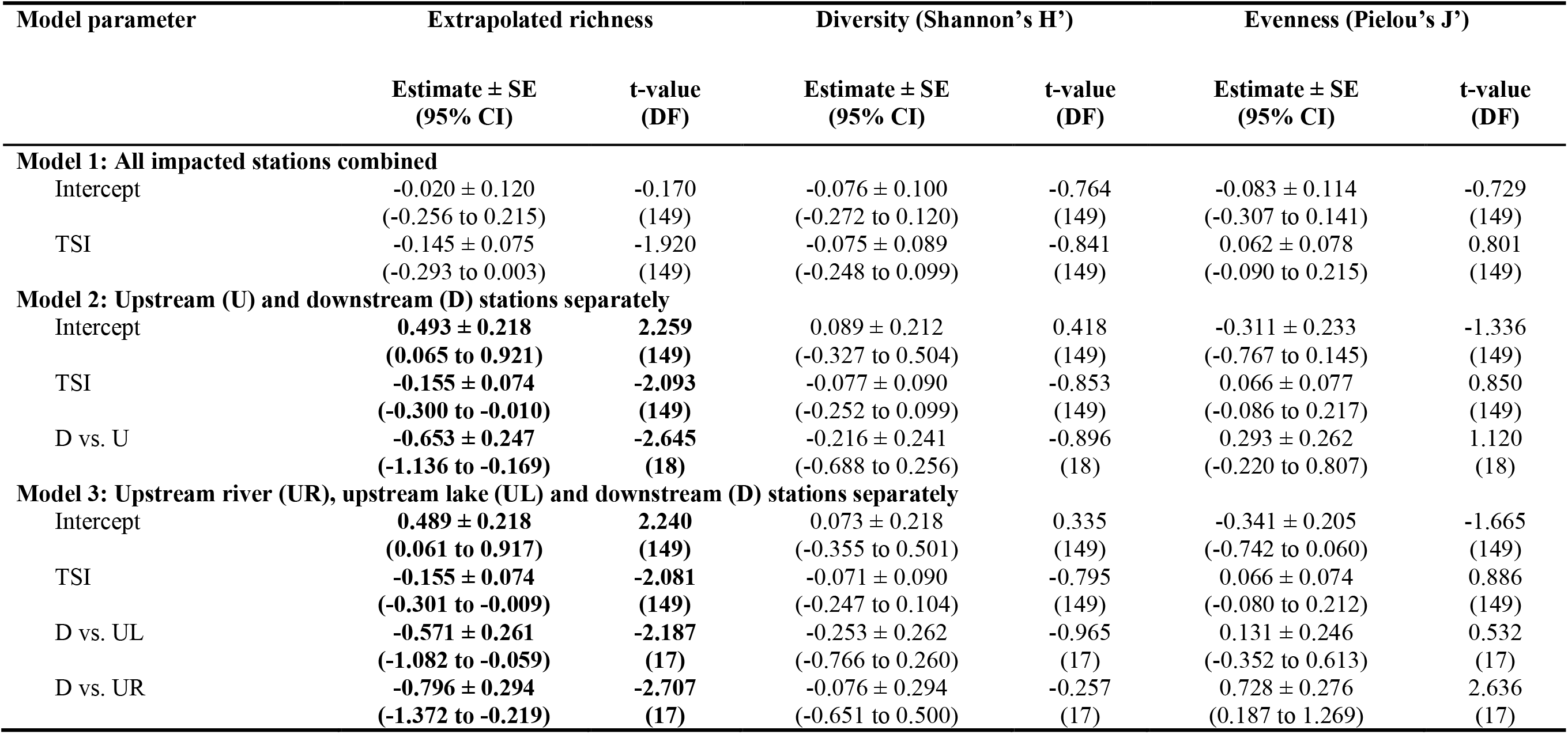
Estimate ± Standard error (SE), 95% Confidence intervals (95% CI), t-values and Degrees of Freedom (DF) of model parameters used to predict change in extrapolated richness (Double jackknife estimation method), diversity (Shannon’s H’) and evenness (Pielou’s J’) in La Grande mega-hydroelectricity complex (3 reservoirs, 22 stations). General linear mixed effects models were used to evaluate the additive effect of time since impoundment (TSI) and categories of impacted stations (D vs. Up, or D vs. UL, vs. UR) on diversity metrics. Predictors that did not include 0 within their 95% CI (*i.e.*, statistically “significant”) are in bold. Downstream stations are used as contrasts in the models.

**Tabel S7.**
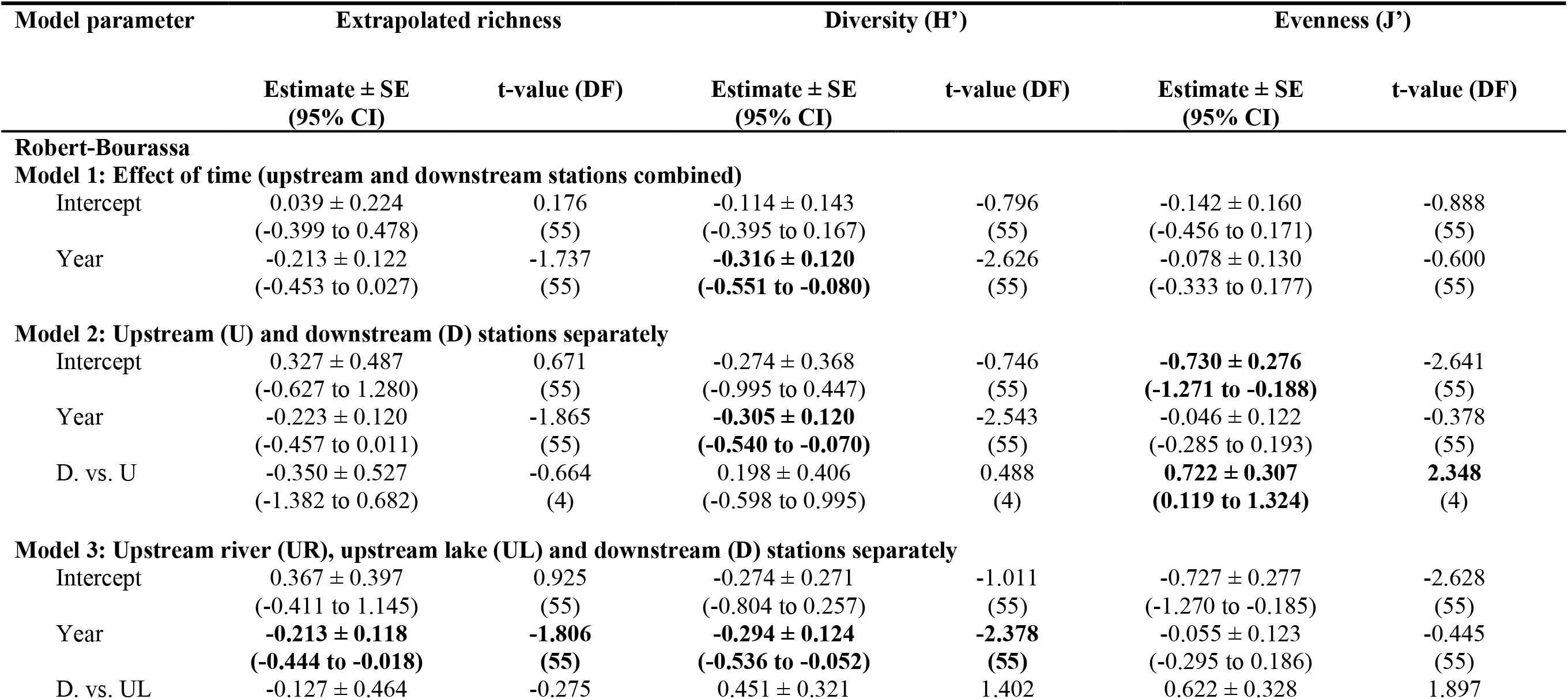
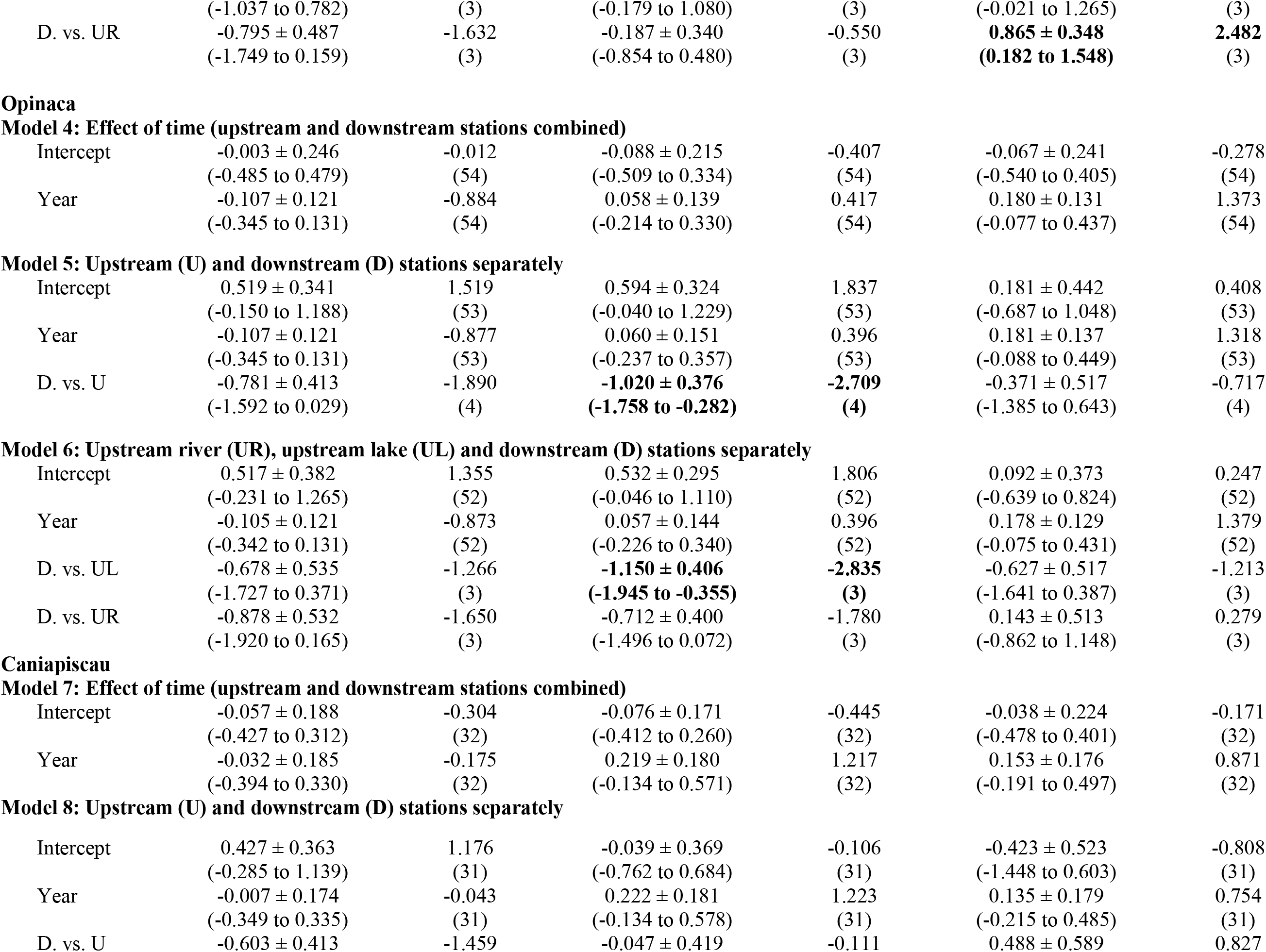
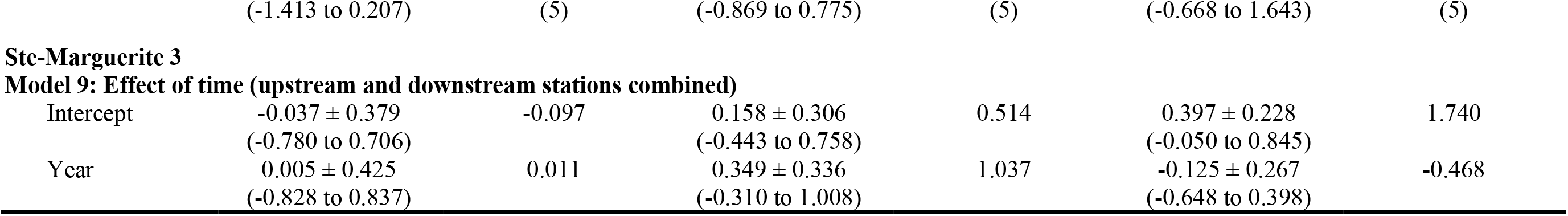
Estimate ± Standard error (SE), 95% Confidence intervals (95% CI) and t-values of model parameters used to predict change in extrapolated richness (Double jackknife estimation), diversity (Shannon-Weaver H’) and evenness (Pielou J’) in La Grande complex reservoirs and Ste-Marguerite 3 reservoirs. General linear mixed effects models were used to evaluate the additive effect of time since impoundment (TSI) and categories of impacted stations (D vs. Up, or D vs. UL, vs. UR) on diversity metrics. Predictors that did not include 0 within their 95% CI (*i.e.*, statistically “significant”) are in bold. Downstream stations are used as contrasts in the models.

**Figure S1.**
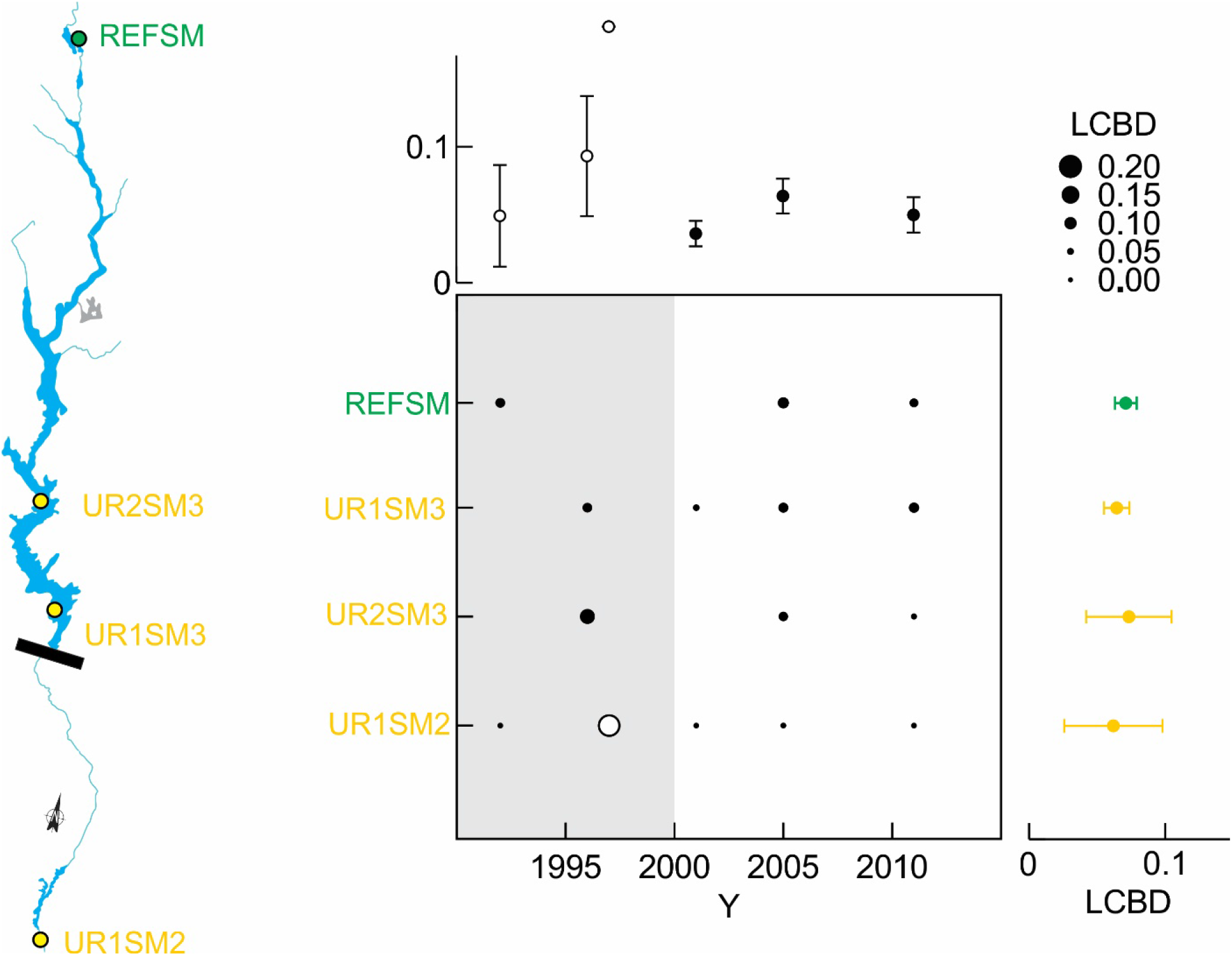
Local contribution to beta-diversity (LCBD) at the reservoir level in SM complex. LCBD values indicate the extent to which each local community is unique in terms of its species composition. Circle surface areas are proportional to the LCBD values. Circles filled in white indicate significant LCBD indices at p>0.05. The upper panel represents mean values of LCBD per year and the right panel represents mean values of LCBD per station. Reference sites are labelled in green, and upstream stations in orange.

**Figure S2.**
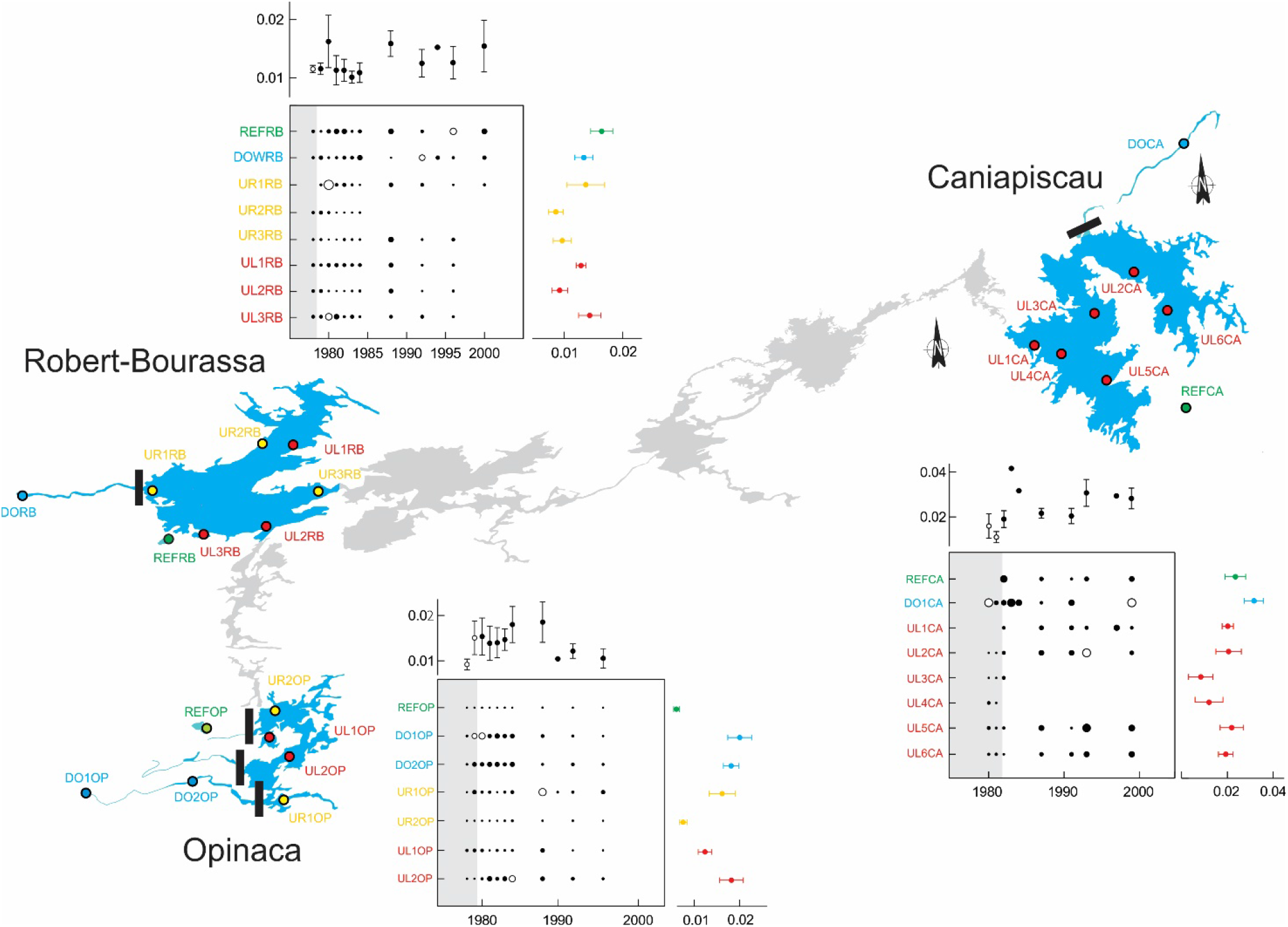
Local contribution to beta-diversity (LCBD) at the reservoir level in LG complex. LCBD values indicate the extent to which each local community is unique in terms of its species composition. Circle surface areas are proportional to the LCBD values. Circles filled in white indicate significant LCBD indices at p<;0.05. The upper panel represents mean values of LCBD per year and the right panel represents mean values of LCBD per station. Reference sites are labelled in green, downstream stations in blue and upstream stations in orange and red. Upstream stations are separated in two categories. Stations with a label starting with “UL” represent stations that were lakes before being a reservoir (orange) and the stations with a label starting with “UR” were rivers or stream before being a reservoir (red).

**Tabel S8.**
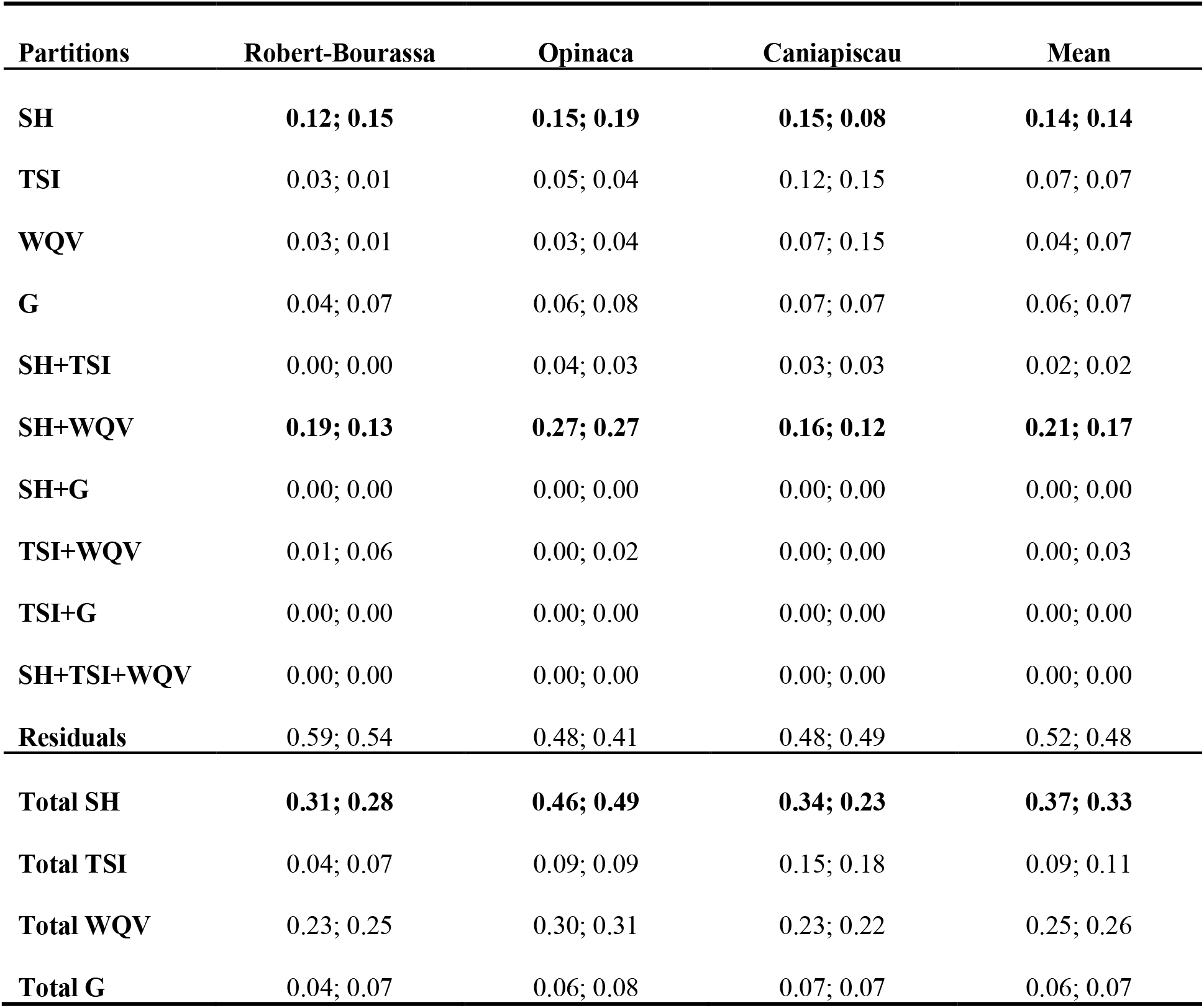
Variation partitioning results at the reservoir level. Variation explained (adjusted R^2^ statistics) by the Spatial heterogeneity [SH] Time since impoundment [TSI], Water quality variables [WQV], Fishing gear [G] matrices, their shared fractions and residual variation. Results are presented for analyses including and excluding reference sites (With; Without). For the significant variables in each matrix, see Fig. 4.

**Tabel S9.**
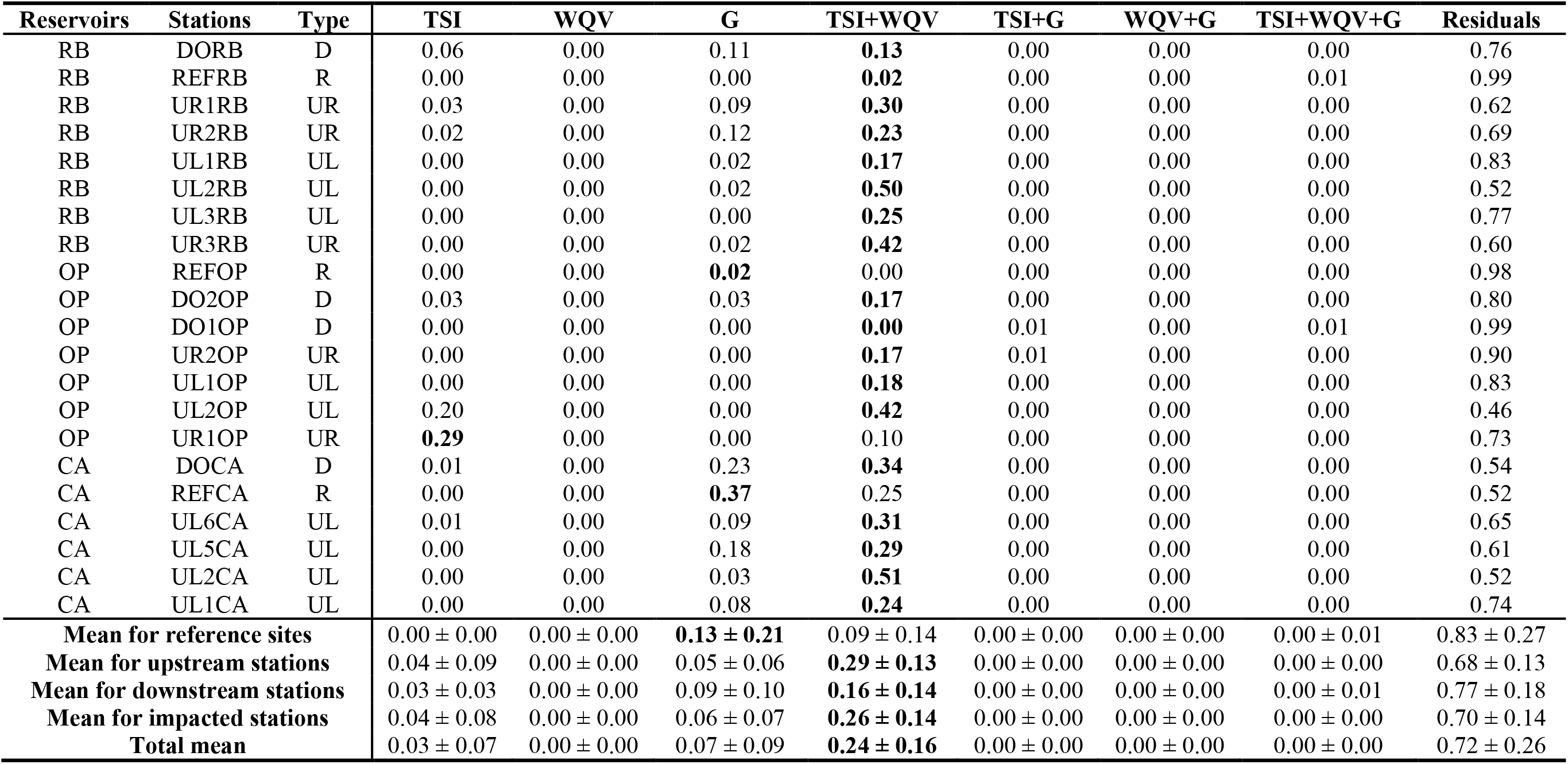
Variation partitioning results at the station level in the LG complex. Variation explained (adjusted R^2^ statistics) by the Spatial heterogeneity [SH] Time since impoundment [TSI], Water quality variables [WQV], Fishing gear [G] matrices, their shared contributions and residual variation. Station are listed by their type (D = downstream, R = reference stations, UR = upstream station that was a river or a stream before impoundment and UL = upstream station that was a lake before impoundment).

**Tabel S10.**
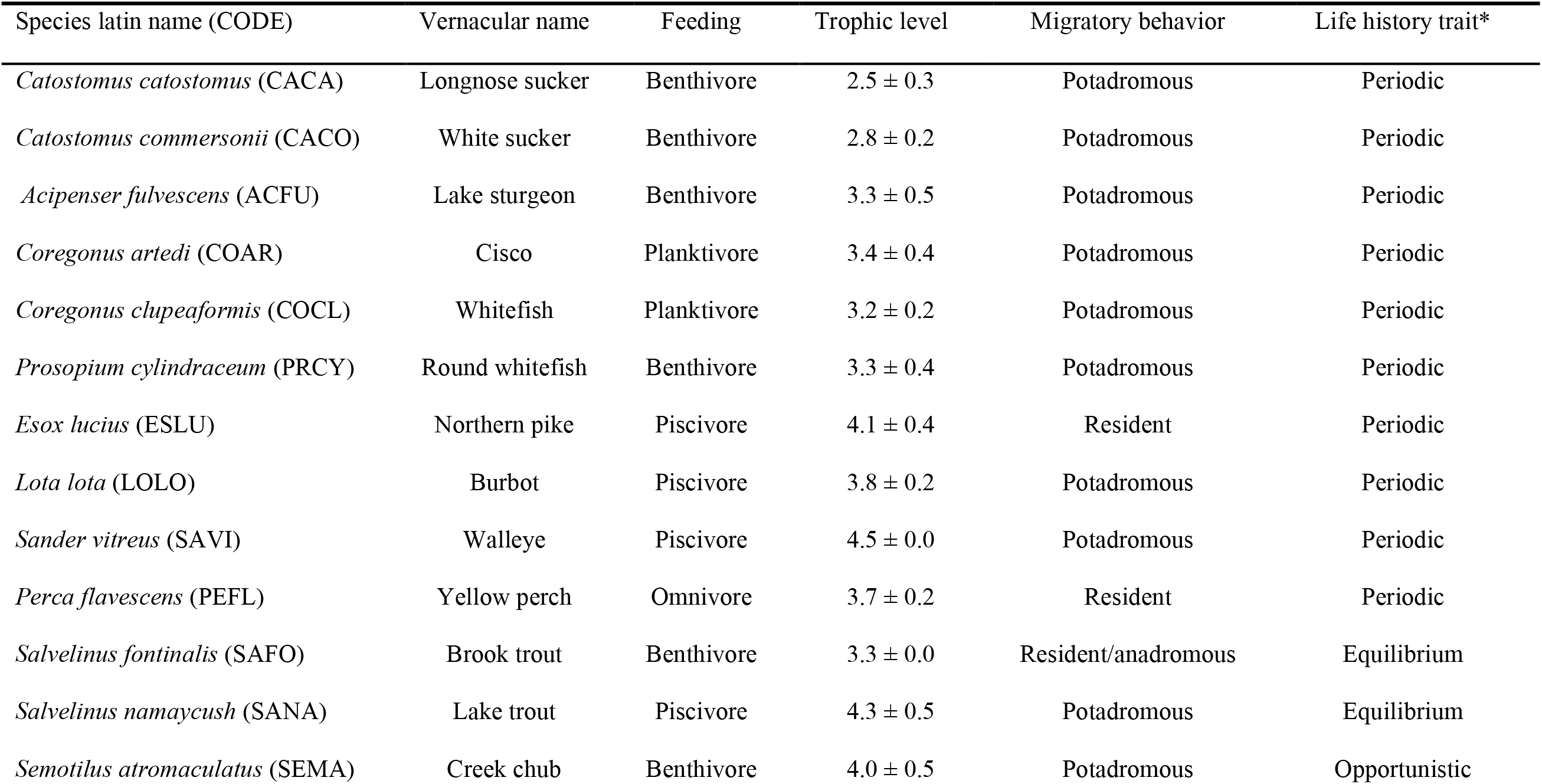
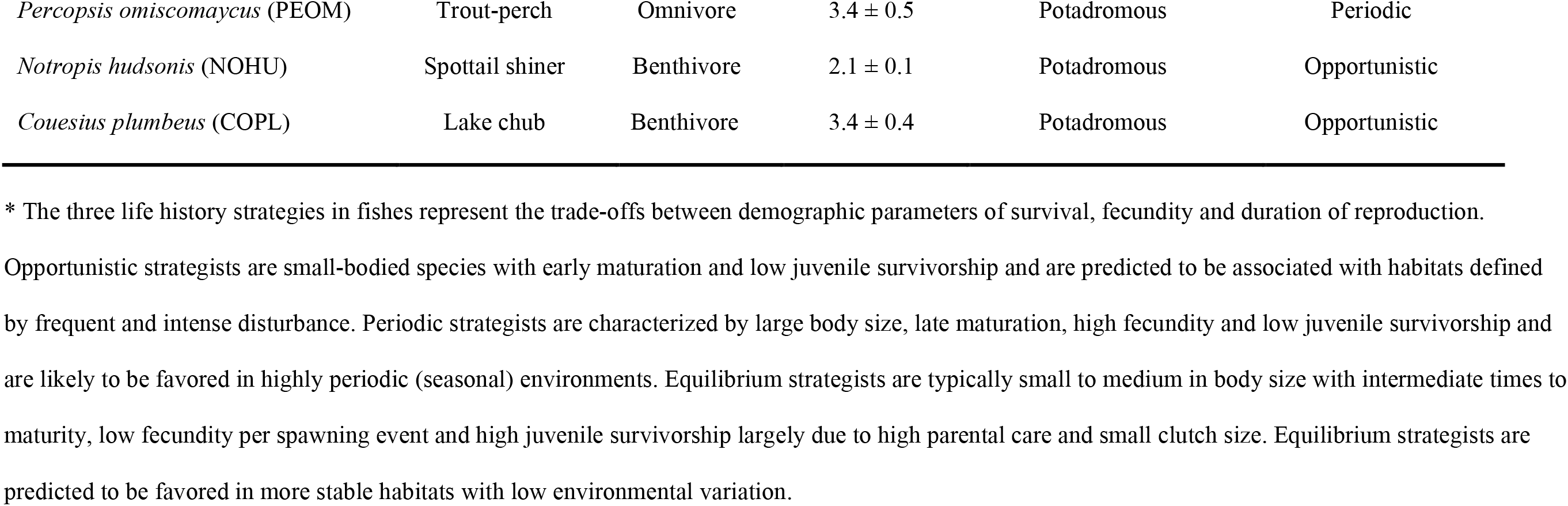
Fish species information. Fish species name (latin name (CODE) and vernacular), feeding habits, trophic level, migratory behavior and life history trait characteristics. Information have been mostly extracted from Fishbase (trophic level and feeding), Olden et al. (2006) and Mims & Olden (2013) (life history trait) and from Desroches and Picard (2013; feeding and migratory behavior).

## Literature Cited

Agostinho, A.A., Julio Jr, H.F. & Petrere Jr., M. (1994) Itaipu Reservoir (Brazil) : Impacts of the Impoundment on the Fish Fauna and Fisheries. Rehabilitation of Freshwater Fisheries, by I.G. Cowx., pp. 171–184. Fishing news Books, Blackwell Scientific Publications, Osney Mead, Oxford, UK.

Agostinho, A.A., Pelicice, F.M. & Gomes, L.C. (2008) Dams and the fish fauna of the Neotropical region: impacts and management related to diversity and fisheries. Brazilian Journal of Biology, 68, 1119–1132.

Blanchet, F.G., Legendre, P. & Borcard, D. (2008) Forward selection of explanatory variables. Ecology, 89, 2623–2632.

Bonner, T.H. & Wilde, G.R. (2000) Changes in the Canadian River Fish Assemblage Associated with Reservoir Construction. Journal of Freshwater Ecology, 15, 189–198.

Burnham, K.P. & Anderson, D.R. (2002) Model Selection and Multimodel Inference: A Practical Information-Theoretic Approach. Second Edition. Springer-Verlag, New York.

Carmignani, J.R. & Roy, A.H. (2017) Ecological impacts of winter water level drawdowns on lake littoral zones: a review. Aquatic Sciences, 1–22.

Clavero, M. & Hermoso, V. (2010) Reservoirs promote the taxonomic homogenization of fish communities within river basins. Biodiversity and Conservation, 20, 41–57.

Deslandes, J.-C. & Fortin, R. (1994) Optimisation Du Réseau de Suivi Environnemental Des Populations de Poissons Des Milieux Affectés Par l’aménagement Du Complexe La Grande (Phase I).

Dudgeon, D., Arthington, A.H., Gessner, M.O., Kawabata, Z.-I., Knowler, D.J., Lévêque, C., Naiman, R.J., Prieur-Richard, A.-H., Soto, D., Stiassny, M.L.J. & Sullivan, C.A. (2006) Freshwater biodiversity: importance, threats, status and conservation challenges. Biological Reviews, 81, 163–182.

Elliott, J.M. (1990) The need for long-term investigations in ecology and the contribution of the Freshwater Biological Association. Freshwater Biology, 23, 1–5.

Fréchette, J.-L. (1980) Réseau Surveillance Écologique. Vol. 1. Société d’énergie de la Baie James. Environnement.

Friedl, G. & Wüest, A. (2002) Disrupting biogeochemical cycles - Consequences of damming. Aquatic Sciences, 64, 55–65.

Galat, D.L., Fredrickson, L.H., Humburg, D.D., Bataille, K.J., Bodie, J.R., Dohrenwend, J., Gelwicks, G.T., Havel, J.E., Helmers, D.L., Hooker, J.B., Jones, J.R., Knowlton, M.F., Kubisiak, J., Mazourek, J., McColpin, A.C., Renken, R.B. & Semlitsch, R.D. (1998) Flooding to Restore Connectivity of Regulated, Large-River Wetlands. BioScience, 48, 721–733.

Gehrke, P.C., Gilligan, D.M. & Barwick, M. (2002) Changes in fish communities of the Shoalhaven River 20 years after construction of Tallowa Dam, Australia. River Research and Applications, 18, 265–286.

Gido, K.B., Guy, C.S., Strakosh, T.R., Bernot, R.J., Hase, K.J. & Shaw, M.A. (2002) Long-Term Changes in the Fish Assemblages of the Big Blue River Basin 40 Years after the Construction of Tuttle Creek Reservoir. Transactions of the Kansas Academy of Science, 105, 193–208.

Gido, K.B., Matthews, W.J. & Wolfinbarger, W.C. (2000) Long-term changes in a reservoir fish assemblage: stability in an unpredictable environment. Ecological Applications, 10, 1517–1529.

Gomes, L. c. & Miranda, L. e. (2001) Riverine characteristics dictate composition of fish assemblages and limit fisheries in reservoirs of the upper Paraná River basin. Regulated Rivers: Research & Management, 17, 67–76.

Grill, G., Lehner, B., Lumsdon, A.E., MacDonald, G.K., Zarfl, C. & Liermann, C.R. (2015) An index-based framework for assessing patterns and trends in river fragmentation and flow regulation by global dams at multiple scales. Environmental Research Letters, 10, 015001.

Grimard, Y. & Jones, H.G. (1982) Trophic Upsurge in New Reservoirs: A Model for Total Phosphorus Concentrations. Canadian Journal of Fisheries and Aquatic Sciences, 39, 1473–1483.

Gubiani, É.A., Gomes, L.C., Agostinho, A.A. & Baumgartner, G. (2010) Variations in fish assemblages in a tributary of the upper Paraná River, Brazil: A comparison between pre and post-closure phases of dams. River Research and Applications, 26, 848–865.

Haxton, T.J. & Findlay, C.S. (2009) Variation in large-bodied fish-community structure and abundance in relation to water-management regime in a large regulated river. Journal of Fish Biology, 74, 2216–2238.

International Energy Agency (IEA). (2016) Key World Energy Statistics. Paris, Cedex, France.

Johnson, P.T., Olden, J.D. & Vander Zanden, M.J. (2008) Dam invaders: impoundments facilitate biological invasions into freshwaters. Frontiers in Ecology and the Environment, 6, 357–363.

Kroger, R.L. (1973) Biological Effects of Fluctuating Water Levels in the Snake River, Grand Teton National Park, Wyoming. American Midland Naturalist, 89, 478–481.

Legendre, P. & De Cáceres, M. (2013) Beta diversity as the variance of community data: dissimilarity coefficients and partitioning. Ecology Letters, 16, 951–963.

Legendre, P. & Gauthier, O. (2014) Statistical methods for temporal and space–time analysis of community composition data. Proceedings of the Royal Society of London B: Biological Sciences, 281, 20132728.

Li, B., Madden, L.V. & Xu, X. (2012) Spatial analysis by distance indices: an alternative local clustering index for studying spatial patterns. Methods in Ecology and Evolution, 3, 368–377.

Liermann, C.R., Nilsson, C., Robertson, J. & Ng, R.Y. (2012) Implications of Dam Obstruction for Global Freshwater Fish Diversity. BioScience, 62, 539–548.

Liew, J.H., Tan, H.H. & Yeo, D.C.J. (2016) Dammed rivers: impoundments facilitate fish invasions. Freshwater Biology, 61, 1421–1429.

Lima, A.C., Agostinho, C.S., Sayanda, D., Pelicice, F.M., Soares, A.M.V.M. & Monaghan, K.A. (2016) The rise and fall of fish diversity in a neotropical river after impoundment. Hydrobiologia, 763, 207–221.

Martinez, P.J., Chart, T.E., Trammell, M.A., Wullschleger, J.G. & Bergersen, E.P. (1994) Fish species composition before and after construction of a main stem reservoir on the White River, Colorado. Environmental Biology of Fishes, 40, 227–239.

McDowall, R.M. (2008) Why are so many boreal freshwater fishes anadromous? Confronting ‘conventional wisdom’. Fish and Fisheries, 9, 208–213.

de Mérona, B. de, Vigouroux, R. & Tejerina-Garro, F.L. (2005) Alteration of Fish Diversity Downstream from Petit-Saut Dam in French Guiana. Implication of Ecological Strategies of Fish Species. Hydrobiologia, 551, 33–47.

Messager, M.L., Lehner, B., Grill, G., Nedeva, I. & Schmitt, O. (2016) Estimating the volume and age of water stored in global lakes using a geo-statistical approach. Nature Communications, 7, 13603.

Nilsson, C., Reidy, C.A., Dynesius, M. & Revenga, C. (2005) Fragmentation and Flow Regulation of the World’s Large River Systems. Science, 308, 405–408.

Okada, E.K., Agostinho, A.A. & Gomes, L.C. (2005) Spatial and temporal gradients in artisanal fisheries of a large Neotropical reservoir, the Itaipu Reservoir, Brazil. Canadian Journal of Fisheries and Aquatic Sciences, 62, 714–724.

Oksanen, J., Blanchet, G., Friendly, M., Kindt, R., Legendre, P., McGlinn, D., Minchin, P.R., O’Hara, R.B., Simpson, G.L., Solymos, P., Stevens, H.H., Szoecs, E. & Wagner, H. (2016) Vegan: Community Ecology Package. R Package Version 2. 4–1.

Pelicice, F.M., Pompeu, P.S. & Agostinho, A.A. (2015) Large reservoirs as ecological barriers to downstream movements of Neotropical migratory fish. Fish and Fisheries, 16, 697–715.

Peres-Neto, P.R., Legendre, P., Dray, S. & Borcard, D. (2006) Variation Partitioning of Species Data Matrices: Estimation and Comparison of Fractions. Ecology, 87, 2614–2625.

Poff, N.L., Olden, J.D., Merritt, D.M. & Pepin, D.M. (2007) Homogenization of regional river dynamics by dams and global biodiversity implications. Proceedings of the National Academy of Sciences, 104, 5732–5737.

Quinn, J.W. & Kwak, T.J. (2003) Fish Assemblage Changes in an Ozark River after Impoundment: A Long-Term Perspective. Transactions of the American Fisheries Society, 132, 110–119.

Reyes-Gavilán, F.G., Garrido, R., Nicieza, A.G., Toledo, M.M. & Braña, F. (1996) Fish community variation along physical gradients in short streams of northern Spain and the disruptive effect of dams. Hydrobiologia, 321, 155–163.

Říha, M., Kubečka, J., Vašek, M., Seda, J., Mrkvička, T., Prchalová, M., Matēna, J., Hladík, M., Čech, M., Draštík, V., Frouzová, J., Hohausová, E., Jarolím, O., Jůza, T., Kratochvíl, M., Peterka, J. & Tušer, M. (2009) Long-term development of fish populations in the Římov Reservoir. Fisheries Management and Ecology, 16, 121–129.

Rodriguez Ruiz, A. (1998) Fish species composition before and after construction of a reservoir on the Guadalete River (SW Spain). Archiv für Hydrobiologie, 142, 353–369.

Rosenberg, D.M., McCully, P. & Pringle, C.M. (2000) Global-Scale Environmental Effects of Hydrological Alterations: Introduction. BioScience, 50, 746–751.

Roy, D. & Messier, D. (1989) A review of the effects of water transfers in the La Grande Hydroelectric Complex (Québec, Canada). Regulated Rivers: Research & Management, 4, 299–316.

Sá-Oliveira, J.C., Hawes, J.E., Isaac-Nahum, V.J. & Peres, C.A. (2015) Upstream and downstream responses of fish assemblages to an eastern Amazonian hydroelectric dam. Freshwater Biology, 60, 2037–2050.

Stoffer, D. (2014) astsa: Applied Statistical Time Series Analysis. R package version 1.3.

Sutela, T. & Vehanen, T. (2008) Effects of water-level regulation on the nearshore fish community in boreal lakes. Hydrobiologia, 613, 13–20.

Taylor, C.A., Knouft, J.H. & Hiland, T.M. (2001) Consequences of stream impoundment on fish communities in a small North American drainage. Regulated Rivers: Research & Management, 17, 687–698.

Tereshchenko, V.G. & Strel’nikov, A.S. (1997) Analysis of long-term changes in the fish community in Rybinsk reservoir. Journal of Ichthyology, 37, 590–598.

Turgeon, K., Solomon, C.T., Nozais, C. & Gregory-Eaves, I. (2016) Do novel ecosystems follow predictable trajectories? Testing the trophic surge hypothesis in reservoirs using fish. Ecosphere, 7, n/a–n/a.

Turgeon, K., Turpin, C. & Gregory-Eaves, I. (in prep.) Do dams affect fish biodiversity in similar ways across biomes? A global quantitative synthesis. in prep.

Vörösmarty, C.J., McIntyre, P.B., Gessner, M.O., Dudgeon, D., Prusevich, A., Green, P., Glidden, S., Bunn, S.E., Sullivan, C.A., Liermann, C.R. & Davies, P.M. (2010) Global threats to human water security and river biodiversity. Nature, 467, 555–561.

Ward, J.V. & Stanford, J.A. (1995) The serial discontinuity concept: Extending the model to floodplain rivers. Regulated Rivers: Research & Management, 10, 159–168.

Whittaker, R.J., Willis, K.J. & Field, R. (2001) Scale and species richness: towards a general, hierarchical theory of species diversity. Journal of Biogeography, 28, 453–470.

Wiens, J.A. (1989) Spatial Scaling in Ecology. Functional Ecology, 3, 385–397.

Winemiller, K.O., McIntyre, P.B., Castello, L., Fluet-Chouinard, E., Giarrizzo, T., Nam, S., Baird, I.G., Darwall, W., Lujan, N.K., Harrison, I., Stiassny, M.L.J., Silvano, R. a. M., Fitzgerald, D.B., Pelicice, F.M., Agostinho, A.A., Gomes, L.C., Albert, J.S., Baran, E., Petrere, M., Zarfl, C., Mulligan, M., Sullivan, J.P., Arantes, C.C., Sousa, L.M., Koning, A.A., Hoeinghaus, D.J., Sabaj, M., Lundberg, J.G., Armbruster, J., Thieme, M.L., Petry, P., Zuanon, J., Vilara, G.T., Snoeks, J., Ou, C., Rainboth, W., Pavanelli, C.S., Akama, A., Soesbergen, A. van & Sáenz, L. (2016) Balancing hydropower and biodiversity in the Amazon, Congo, and Mekong. Science, 351, 128–129.

Zarfl, C., Lumsdon, A.E., Berlekamp, J., Tydecks, L. & Tockner, K. (2014) A global boom in hydropower dam construction. Aquatic Sciences, 77, 161–170.

## References

Froese, R. and D. Pauly. Editors. 2016. FishBase. World Wide Web electronic publication. www.fishbase.org, version 10.2016

Desroches, J.-F. and Picard, I. (2013). Poissons d’eau douce du Québec et des Maritimes. Éditions Michel Quintin. Québec. Canada

Mims M.C. & Olden J.D. (2013) Fish assemblages respond to altered flow regimes via ecological filtering of life history strategies. Freshwater Biology 58, 50–62.

Olden J.D., Poff, N.L. & Bestgen K.R. (2006) Life-history strategies predict fish invasions and extirpations in the Colorado River basin. Ecological Monographs 76(1), 25–40.

